# Activity-dependent Mitochondrial ROS Signaling Regulates Recruitment of Glutamate Receptors to Synapses

**DOI:** 10.1101/2023.08.07.552290

**Authors:** Rachel L. Doser, Kaz M. Knight, Ennis Deihl, Frederic J. Hoerndli

## Abstract

Our understanding of mitochondrial signaling in the nervous system has been limited by the technical challenge of analyzing mitochondrial function *in vivo*. In the transparent genetic model *Caenorhabditis elegans,* we were able to manipulate and measure mitochondrial signaling as well as neuronal activity *in vivo*. Using this approach, we provide evidence supporting a novel role for mitochondrial signaling in dendrites. Specifically, we show that dendritic mitochondria take up calcium (Ca^2+^) via the mitochondrial Ca^2+^ uniporter MCU-1 causing an upregulation of mitochondrial reactive oxygen species (mitoROS) production. We also observed that mitochondria are positioned in close proximity to synaptic clusters of GLR-1, the *C. elegans* ortholog of the AMPA subtype of glutamate receptors that mediate neuronal excitation. We show that synaptic recruitment of GLR-1 is downregulated by mitoROS signaling resulting from mitochondrial Ca^2+^ uptake via MCU-1. Thus, postsynaptic mitochondria provide a means of negative feedback that regulates excitatory synapse function which may be vital for neuronal homeostasis by preventing excitotoxicity and energy depletion.

## INTRODUCTION

As the predominant excitatory synapse type in the brain, glutamatergic synapses are important for organismal physiology and homeostasis as well as much of the brain’s processing^1–3^. Plasticity, or the change in efficacy, of these synapses underlies adaption to environmental conditions as well as learning and memory formation. Although presynaptic changes contribute to synaptic strength, the number of ionotropic glutamate receptors, especially the *α*-amino-3-hydroxy-5-methyl-4-isoxazole (AMPA) subtype (AMPARs), at the postsynaptic membrane is a strong correlate of synaptic strength. Changes in synaptic expression of AMPARs is a calcium (Ca^2+^)-dependent, multi-step process involving long-distance transport of the receptors by molecular motors^4–9^, delivery of AMPAR-containing vesicles to synaptic sites^10, 11^, exocytosis and endocytosis of AMPARs to the membrane^12, 13^ as well as reorganization of postsynaptic proteins and cytoskeletal architecture^14–16^.

The mechanisms underlying postsynaptic plasticity are metabolically demanding processes requiring the upregulation of mitochondrial metabolism to meet energy demands^17, 18^. There is growing evidence that mitochondria are also important for other cellular functions including regulation of gene expression, Ca^2+^ homeostasis, inflammatory signaling, and lipid biogenesis^19, 20^. Interestingly, the generation of reactive oxygen species (ROS), such as superoxide and hydrogen peroxide, by the mitochondrial respiratory chain and other matrix proteins^21^ is gaining traction as an essential signaling mechanism with many identified downstream effectors in neurons^22, 23^. It has become clear that ROS act as a physiological signal^22^ that are necessary for neuronal development^24^, excitatory and inhibitory neurotransmission^25^, as well as synaptic plasticity^26, 27^.

For instance, evidence accumulated over the last 25 years has demonstrated that ROS signaling is required for normal synaptic expression of AMPARs. Early evidence came from results suggesting abnormal plasticity of glutamatergic synapses, learning and memory when ROS are elevated or diminished^26, 28–30^. Since these studies, we and others have shown that ROS signaling can regulate the number of synaptic AMPARs via ROS-dependent regulation of AMPAR phosphorylation^31^ or the long-distance transport and delivery of AMPARs to synapses^32, 33^. Despite our understanding of several downstream effectors of ROS signaling, it is unclear when or where ROS signaling originates in neurons. As previously mentioned, ROS is predominantly generated as a byproduct of mitochondrial respiration but is also produced by NADPH oxidase and peroxisome enzymes^22^. Despite mitochondria being the major source of ROS, it has not been assessed *in vivo* if or how mitochondrial ROS (mitoROS) production is regulated by neuronal activity. In addition, mitochondria are positioned at pre- and postsynaptic sites^34^ where they likely contribute to synaptic function. However, our understanding of the roles mitochondria play at synapses has been limited by our ability to study mitochondrial function *in vivo* under physiological conditions.

The transparent nematode *Caenorhabditis elegans* is a powerful genetic model that has been widely accepted for studying mitochondrial function, Ca^2+^ handling and ROS signaling *in vivo* especially in the context of aging and neurodegeneration^35–39^. Additionally, *C. elegans* have been used extensively in neuroscience research^40^ due to their relatively simple nervous system composed of neurons whose gene expression and synaptic connections are completely mapped^41, 42^. Importantly, most of the key players at glutamatergic synapses are conserved, including subunits of AMPARs and other glutamate receptor subtypes^43^, and are regulated in similar fashion to their vertebrate orthologues^8, 11, 44, 45^. Using *C. elegans* to study the regulation of glutamatergic synapses, we have shown that Ca^2+^ signaling regulates transport and delivery of GLR-1, the *C. elegans* ortholog of the AMPAR subunit GluA1, to synapses. Moreover, our previous work revealed that ROS signaling interacts with Ca^2+^ signaling in the cell body to control the amount of GLR-1 transport as well as in dendrites to regulate synaptic delivery of GLR-1^32^. Thus, an interplay between ROS and Ca^2+^ signaling at postsynaptic sites appears to be important for AMPAR localization to synapses, but the role of postsynaptic mitochondria in this process has not been addressed.

Here, using *in vivo* imaging and optogenetic tools in *C. elegans,* we assessed the role of postsynaptic mitochondria as signaling hubs that integrate neuronal activity and regulate AMPAR localization to synapses. We found that in response to neuronal activation, mitochondria take up Ca^2+^ causing an increase in their ROS production. Most dendritic mitochondria were located in close proximity to clusters of surface localized GLR-1, which are representative of postsynaptic sites. To demonstrate functional relevance of activity dependent mitoROS signaling, we show that activity-dependent mitoROS production, requiring the mitochondrial Ca^2+^ uniporter MCU-1 regulates transport, delivery and recruitment of GLR-1 to synapses. Since the number of glutamate receptors at a synapse controls the efficacy of excitatory transmission, activity-induced mitoROS production may constitute a critical inhibitory feedback mechanism that balances neuronal excitability with cellular energy capacity.

## RESULTS

### Activity-dependent Mitochondrial Ca^2+^ Uptake Regulates Synaptic Recruitment of GLR-1

As in vertebrates, the majority of neuronal activation in *C. elegans* is due to glutamatergic transmission. Activation occurs when glutamate is released from a presynaptic neuron which binds to and opens the cation pore of postsynaptic glutamate receptors, including AMPARs. Influx of cations into the postsynaptic neuron initiates opening of voltage gated Ca^2+^ channels which causes a rapid increase in cytoplasmic Ca^2+^. This Ca^2+^ activates a multitude of signaling cascades before being rapidly taken up by the endoplasmic reticulum and mitochondria or extruded to extracellular space^46^. Mitochondria in various neuronal subtypes have discrete Ca^2+^ handling capabilities^47^, so we first characterized mitochondrial Ca^2+^ uptake *in vivo* in the neurites of the AVA glutamatergic interneurons. To do this, we co-expressed the light-sensitive cation channel ChRimson ^48^ with the mitochondrial calcium indicator mitoGCaMP^49^ targeted to the inner mitochondrial matrix (Figure 1A). This combination of tools allowed us to measure Ca^2+^ uptake by individual mitochondria following repetitive optical activation. It is important to note that our photoactivation protocol involved optical stimulation at 33.3 mHz, a rate that is similar to the spontaneous activity of AVA neurons^32^. This assay revealed that there is diversity in Ca^2+^ handling among dendritic mitochondria. Some mitochondria take up the most Ca^2+^ upon the first optical activation (Mito 1; Figure 1B and 1C) whereas others uptake more Ca^2+^ following the second or third stimulation (Mito 2; Figure 1B and 1C).

**Figure 1:**
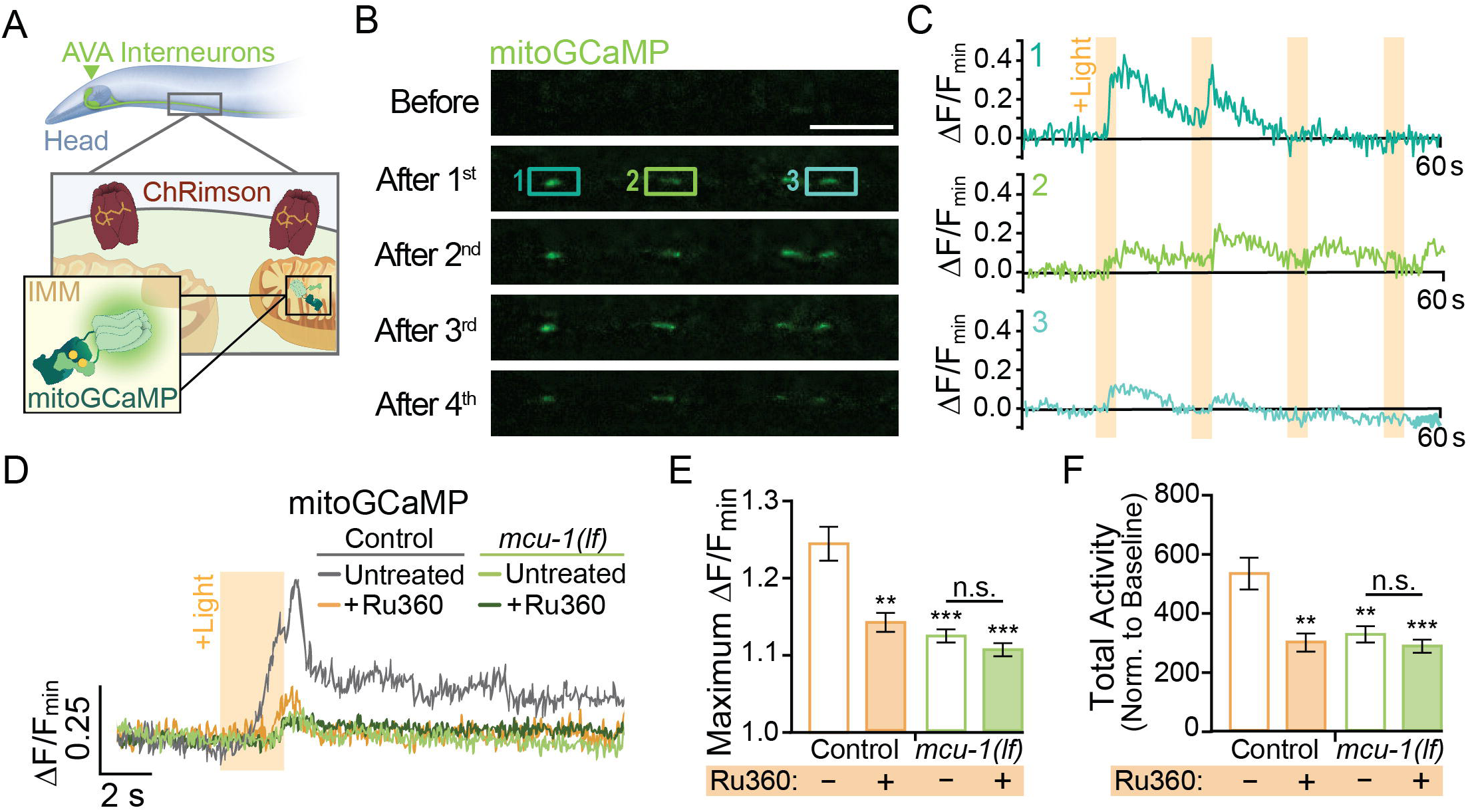
Neuronal activity causes mitochondrial Ca^2+^ uptake via MCU-1. A) Illustration depicting transgenic expression and subcellular location of ChRimson and mitoGCaMP in the AVA neurons. B) Representative images of mitoGCaMP fluorescence in a single z-plane before and after four optical activations. C) Normalized mitoGCaMP fluorescence for the regions of interest in B during repetitive optical activation (+Light, 5 µW at 33.3 mHz). D) Representative normalized mitoGCaMP traces (30 s) following optical stimulation (+Light) of the AVA neurons in worms pre-treated with Ru360 and in untreated controls or *mcu-1(lf)*. E) Normalized amplitude of light-evoked mitoGCaMP events and F) total mitoGCaMP activity (n≥20 mitochondria from 5-8 animals per group). All scale bars = 5 µm. Data is represented as mean ± s.e.m.; n.s. = not significant, **: p<0.005, ***: p<0.0005 compared to controls using a One-Way ANOVA.

Ca^2+^ entry into the matrix is gated by the Ca^2+^-sensitive mitochondrial uniporter MCU-1^50^ which is encoded by the *mcu-1* gene in *C. elegans.* We characterized the effect of the *mcu-1(ju1154)* loss of function allele^51^ (hereafter called *mcu-1(lf)*) on activity-dependent mitochondrial Ca^2+^ uptake by imaging mitoGCaMP in *mcu-1(lf)* (Figure 1D-1F). We found that the amplitude of evoked mitoGCaMP events in *mcu-1(lf)* was drastically decreased compared to controls (Figure 1D-1F). Additionally, the total mitoGCaMP activity, a combined measure of the amplitude and duration of all Ca^2+^ events, was also reduced in *mcu-1(lf)* (Figure 1F). Due to the possibility of functional compensation in *mcu-1(lf)*, we also tested how acute treatment with the Ruthenium compound Ru360, an MCU-1 blocker^52^, alters activity-dependent mitochondrial Ca^2+^ uptake. Following a 10-minute treatment with Ru360, we observed a decrease in the amplitude and total activity of evoked mitoGCaMP events that were similar to *mcu-1(lf).* To test the specificity of Ru360 for inhibiting Ca^2+^ uptake via MCU-1, we treated *mcu-1(lf)* with Ru360 but did not detect additional inhibition of mitochondrial Ca^2+^ uptake (Figure 1D-1F). This Ru360 treatment suppressed mitochondrial Ca^2+^ uptake out to 60 minutes post-treatment (Supplemental Figure 1). This experiment showed that loss or inhibition of MCU-1 almost completely prevents activity-dependent mitochondrial Ca^2+^ uptake.

While imaging mitochondrial-localized fluorescent indicators in the AVA glutamatergic interneurons, we observed that mitochondria are in close proximity to clusters of surface-localized GLR-1, indicative of postsynaptic sites, that were visualized using GLR-1 tagged with pH-senstive GFP (SuperEcliptic pHlourin, SEP) on the N-terminal (Figure 2A). The regulation of mitochondrial function and signaling by Ca^2+^ appears to be integral to synaptic function and plasticity^49, 53–56^ which led us to test if postsynaptic mitochondrial Ca^2+^ uptake is required for normal GLR-1 localization to synapses. First, we quantified SEP::GLR-1 fluorescence in AVA dendrites *in vivo* to assess if the number of GLR-1 at synapses is altered by loss or inhibition of MCU-1. Initial observations revealed a dramatic increase in SEP::GLR-1 in *mcu-1(lf)* mutants (Supplemental Figure 2A) suggesting more synaptic GLR-1. Acute Ru360 treatment slightly, but not significantly, increased the fluorescence of SEP::GLR-1 puncta along the AVA neurite (Supplemental Figure 2A). Next, we used fluorescence recovery after photobleaching (FRAP) of SEP::GLR-1 to measure the rate of GLR-1 recruitment to the synaptic membrane (Supplemental Figure 2B). SEP will only fluoresce when GLR-1 is positioned at the plasma membrane and is quenched while in transport vesicles or synaptic endosomes. Thus, the recovery of SEP fluorescence in a photobleached neurite is a measure of GLR-1 that has been exocytosed to the membrane (Supplemental Figure 2B). The rate of SEP fluorescence recovery (without individual normalization; see Methods for analysis details) was increased more than two-fold in *mcu-1(lf)* and slightly increased following Ru360 treatment (Figure 2B and 2C). When the fluorescence at each timepoint after photobleaching is normalized to the fluorescence before photobleaching, the relative FRAP is unchanged between experimental groups (Supplemental Figure 2C). Taken together, these analyses show that loss or inhibition of MCU-1 leads to excessive recruitment of GLR-1 to synapses, but the rate of exocytosis is increased proportional to the amount of GLR-1 at synapses.

**Figure 2:**
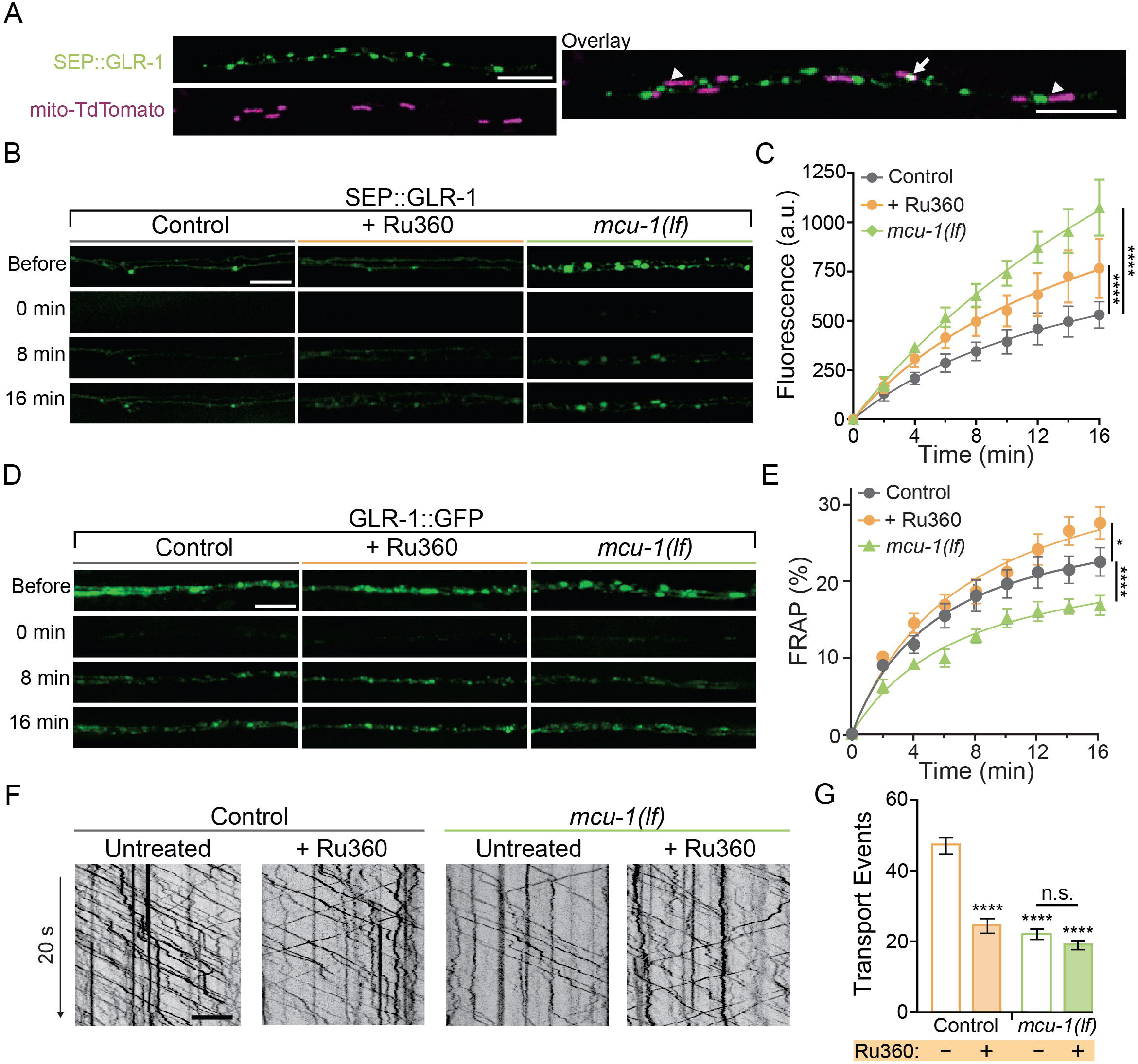
Decreased mitochondrial Ca^2+^ uptake affects transport and recruitment of GLR-1 to synapses. A) Single z-plane fluorescent images of mitochondria (mito-TdTomato) and surface localized GLR-1 (SEP::GLR-1) showing mitochondria localized at (arrows) or adjacent to (arrow heads) SEP::GLR-1 puncta. B and D) Representative images of SEP::GLR-1 (B) or GLR-1::GFP (D) fluorescence before, immediately after, 8-, and 16-minutes post-photobleach (PB). C) Fluorescence (arbitrary units = a.u.) of SEP over 16 minutes post PB (n≥6 animals per group). E) Percent GFP fluorescence recovery after PB (FRAP) over 16 minutes (n≥8 animals per group). *: p<0.01, ****: p<0.0001 using an extra sum-of-squares F-test with a Bonferroni correction. F) 20-second representative kymographs of GLR-1::GFP movement in AVA neurite in controls and *mcu-1(lf)* with or without Ru360 pre-treatment. Time is represented on the y-axis and distance on the x-axis. G) Total transport events quantified from kymographs in all conditions (n>10 animals per group). All scale bars = 5 µm. Data is represented as mean ± s.e.m.; n.s = not significant, ****: p<0.0001 compared to controls or indicated experimental group using a One-Way ANOVA.

The exocytosis rate of GLR-1 depends on the local GLR-1 reserves in synaptic endosomes^16^. Resupplying of these local receptor pools occurs when GLR-1-containing transport vesicles are delivered to endosomes or other local reserves^57^. The delivery rate of new GLR-1 can be measured by FRAP of GLR-1::GFP (Supplemental Figure 2D). In *mcu-1(lf)*, the rate of GLR-1::GFP FRAP in *mcu-1(lf)* was comparable to controls whereas the rate of FRAP in Ru360 treated animals was slightly increased (Figure 2D and 2E). Synaptic delivery of GLR-1 is dependent upon the transport of GLR-1-containing vesicles by molecular motors from the cell body where GLR-1 is predominantly synthesized. Since both *mcu-1(lf)* and Ru360 treatment globally reduce mitochondrial Ca^2+^ uptake, we tested if GLR-1 transport out of the cell body is also sensitive to loss or inhibition of MCU-1 as it would impact downstream GLR-1 delivery and exocytosis rates. We visualized individual GLR-1::GFP transport by photobleaching a section (∼40 µm) of the AVA neurites as previously described^9, 32^. Interestingly, we found that both *mcu-1(lf)* and Ru360 treatment decreased the amount of GLR-1 transport by ∼50% (Figure 2F and 2G). Ru360 treatment of *mcu-1(lf)* did not further decrease the amount of GLR-1 transport compared to *mcu-1(lf)* alone. Together, these results suggest that mitochondrial Ca^2+^ uptake differentially regulates GLR-1 transport out of the cell body and synaptic recruitment of GLR-1.

Previous work has shown that cytoplasmic Ca^2+^ signaling regulates transport and synaptic localization of GLR-1^8, 11, 32^, so we tested if decreased Ca^2+^ uptake alters the amplitude or duration of cytoplasmic Ca^2+^ transients in dendrites following neuronal activation since this would impact downstream Ca^2+^ signaling and synaptic recruitment of GLR-1. We expressed ChRimson and the cytoplasmic Ca^2+^ indicator GCaMP6f in the AVA neurons in *mcu-1(lf)* and control animals (Supplemental Figure 2E). This approach bypasses activation by presynaptic inputs allowing direct activation of the AVA interneurons. We simultaneously optically activated the AVA neurons and recorded GCaMP6f fluorescence in *mcu-1(lf)* and Ru360-treated controls in the same dendritic region of the AVA neurons where GLR-1 transport and FRAP were analyzed. There were no detectable changes in cytoplasmic Ca^2+^ transients in dendrites following AVA activation with ChRimson between *mcu-1(lf)* or with Ru360 treatment compared to controls (Supplemental Figure 2F-2H) suggesting that loss or inhibition of MCU-1 does not drastically alter activity-dependent cytoplasmic Ca^2+^ influx or the duration of a Ca^2+^ event in dendrites. In other words, the loss or inhibition of MCU-1 does not seem to impact synaptic localization of GLR-1 by indirectly modulating cytoplasmic Ca^2+^ signaling.

### Neuronal Excitation Upregulates Mitochondrial ROS Signaling

Our previous work has shown that ROS regulate transport and synaptic delivery of GLR-1^32, 33^. To further address the mechanism by which Ca^2+^ influx by MCU-1 modulates GLR-1, we tested if activity-dependent Ca^2+^ uptake regulates mitochondrial ROS (mitoROS) production. To do this, we stimulated the AVA neuron with ChRimson using the same optical activation that initiated mitochondrial Ca^2+^ uptake (Figure 1). Then, we measured ROS levels at dendritic mitochondria using a genetically encoded ratiometric ROS sensor that was localized to the outer mitochondrial membrane (mito-roGFP^58^; Figure 3A). We found that the duration of repetitive AVA stimulation positively correlated with the mito-roGFP fluorescence ratio (F_ratio_; 405/488 nm) indicating increased ROS following neuronal activation. The F_ratio_ was unchanged in controls that were not treated with retinal, which is required for optical stimulation, and subjected to the light stimulation protocol (Figures 3B and 3C). Similar to mitoGCaMP responses, we saw diversity among dendritic mitochondria in mito-roGFP F_ratios_ in following neuronal activation of the AVA neurons (Supplemental Figure 3A and 3B). The frequency distribution of mito-roGFP F_ratios_ of individual mitochondria without stimulation is unimodal (centered at 0.03) but becomes bimodal following 60 minutes of repetitive activation. One peak is slightly right shifted (centered at 0.05) and the other is strongly right shifted, corresponding to significantly higher mito-roGFP F_ratios_ (centered at 0.09; Supplemental Figure 3A and 3B). These results suggest that mitochondria within these neurites differentially respond to activity in terms of their ROS production.

**Figure 3:**
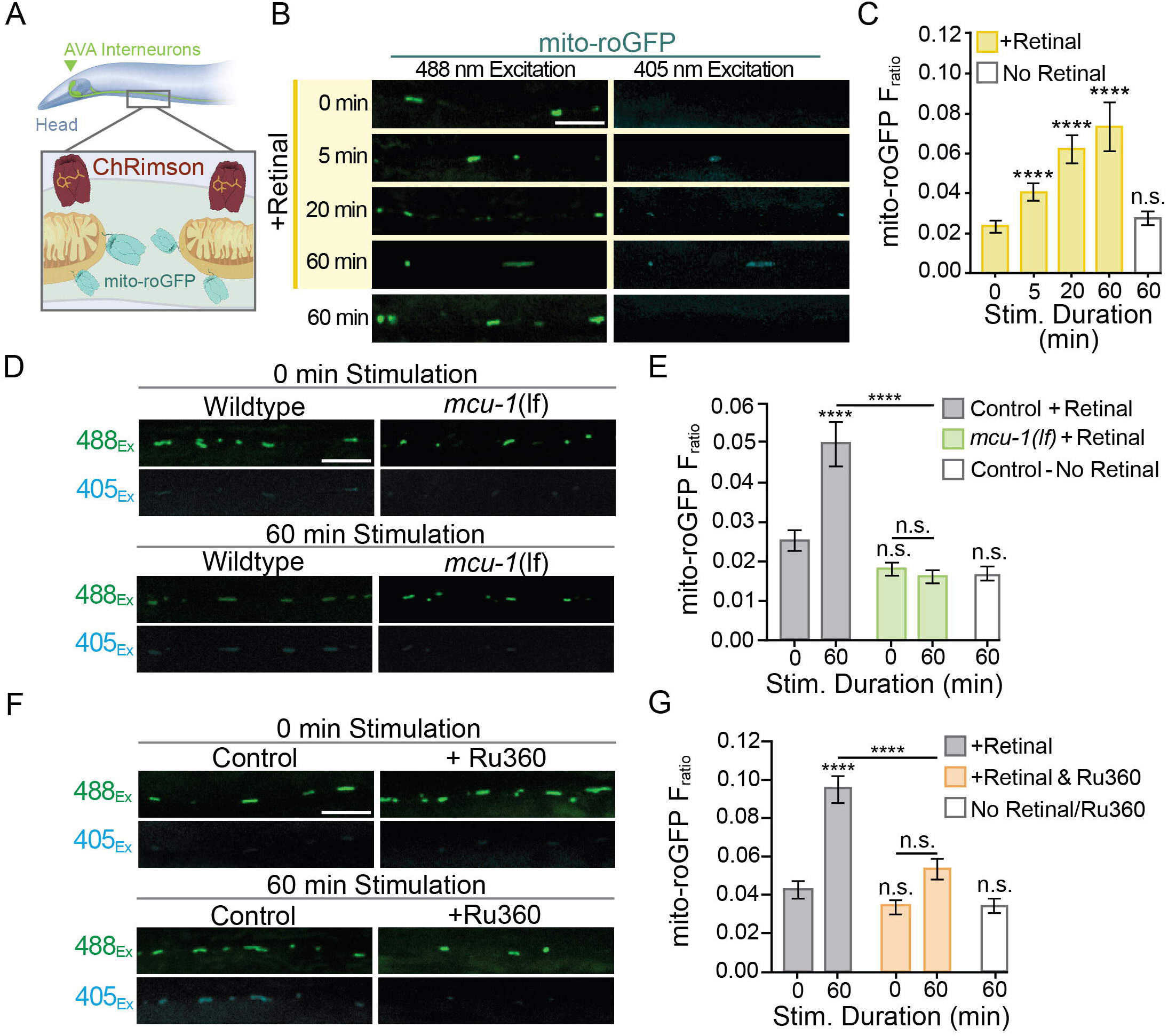
MitoROS production is upregulated by neuronal activity and dependent on mitochondrial Ca^2+^ uptake via MCU-1. A) Illustration showing transgenic expression and subcellular localization of ChRimson and mito-roGFP in the AVA neurons. B) Representative images of mito-roGFP fluorescence in a single z-plane when excited with 488 nm or 405 nm light following optogenetic stimulation with or without all-trans-Retinal. C) Mito-roGFP fluorescence ratio (405/488 nm) following 0, 5, 20 or 60 minutes of repetitive optical stimulation (40 μW/mm^2^ at 33.3 mHz) with Retinal pre-treatment and 60 minutes of repetitive optical stimulation without Retinal pretreatment (n>30 mitochondria from 8 animals per group). D and F) Representative images of mito-roGFP fluorescence in a single z-plane when excited by 488 nm or 405 nm light following 0 or 60 minutes of repetitive optical stimulation with Retinal pre-treatment. E) Mito-roGFP F_ratio_ following 0 or 60 minutes of repetitive optical stimulation in *mcu-1(lf)* and controls as well as non-Retinal treated controls that underwent 60-minutes of stimulation (n≥32 mitochondria from 8 animals per group). Statistical comparisons are between groups and the 0-minute control unless indicated by horizontal bar. G) Mito-roGFP fluorescence ratio at 0 and 60 minutes following repeated optical stimulation with or without Ru360 treatment (n≥38 mitochondria from 8 animals per group). All scale bars = 5 µm. Data is represented as mean ± s.e.m.; n.s. = not significant, ****: p<0.0001 compared to controls or indicated experimental group using a One-Way ANOVA.

Since optical activation is artificial and does not rely on synaptic transmission, it is possible that mitoROS production is not upregulated by natural neuronal activation. To address this, we took advantage of the well-defined circuitry in *C. elegans* and designed an experiment to activate a subset of mechanosensory neurons that detect physical touch and vibration^59^ and provide excitatory input to AVA neurons. This involved repetitively activating presynaptic mechanosensory neurons with vibration caused by dropping culture plates containing freely behaving worms from a short distance (∼ 5 cm) onto the bench top every 30 seconds for a duration of 5 or 10 minutes. Then, worms were mounted for imaging to assess the F_ratio_ of mito-roGFP. The mito-roGFP F_ratio_ was slightly increased following 5 minutes and significantly increased by 10 minutes of repetitive mechano-stimulation (Supplemental Figure 3C and 3D) indicating that mitoROS production is also increased by native means of neuronal activation. So, does this activity dependent upregulation of mitochondrial ROS production require Ca^2+^ uptake through MCU-1? Expression of ChRimson and mito-roGFP in *mcu-1(lf)* revealed that the loss of MCU-1 prevented activity-induced increases in the mito-roGFP F_ratio_ even after 60 minutes of repetitive optical activation (Figure 3D and 3E; Supplemental Figure 3E). Pre-treatment with Ru360 prior to optical activation similarly prevented the activity-induced increase in mito-roGFP F_ratio_ (Figure 3F and 3G; Supplemental Figure 3F). In summary, both the acute pharmacological inhibition and genetic loss of MCU-1 prevented activity-dependent upregulation of mitoROS production. Furthermore, this activity-induced mitoROS production suggests that mitochondria play an active signaling role during neuronal activation.

### Mitochondrial ROS Signaling Regulates Synaptic Recruitment of GLR-1

We next addressed the possible role of ROS production at dendritic mitochondria in regulating the multistep process required for synaptic recruitment of GLR-1. To acutely induce ROS production at dendritic mitochondria, we expressed the photosensitizer KillerRed which produces ROS (specifically hydrogen peroxide) upon photoactivation (PA) with green light. We localized KillerRed to mitochondria (mitoKR) by anchoring it to the outer mitochondrial membrane with the localization tag TOMM20^60^. First, we co-expressed mitoKR with mito-roGFP (Supplemental Figure 4A and 4B) for optimization of a PA protocol to artificially induce mitoROS production at a subset of synapses (local) or throughout the AVA neuron (global). To test our local PA protocol, we used a microscopy set-up that was equipped for targeted illumination (see Materials and Methods) allowing us to direct a green LED to a small portion (∼10 µm) of the AVA neurites containing 1-3 mitochondria for 15 or 30 seconds (Supplemental Figure 4C). The mito-roGFP F_ratio_ was significantly increased in the mitochondria that were targeted for 15 or 30 seconds of PA when compared to non-activated controls as well as neighboring mitochondria not targeted for PA (Supplemental Figure 4D). This enabled us to locally activate mitoKR (15 s duration) prior to assessing GLR-1 exocytosis via FRAP of SEP::GLR-1 (Figure 4A). Interestingly, local PA dramatically decreased SEP::GLR-1 FRAP in mitoKR-expressing worms compared to controls (Figure 4B and 4C). The FRAP rate of non-activated mitoKR worms was significantly decreased compared to controls, but to a lesser extent than with PA (Supplemental Figure 5A). This is likely due to activation of mitoKR during imaging of SEP fluorescence. This dramatic downregulation of GLR-1 exocytosis due to localized artificial mitoROS production could be caused by altered delivery of GLR-1-containing transport vesicles. When we assessed GLR-1 delivery via FRAP of GLR-1::GFP following local PA of mitoKR, we observed that PA of mitoKR decreased the rate of GLR-1::GFP FRAP in worms expressing mitoKR in comparison to controls lacking mitoKR (Figure 4D and 4E) as well as mitoKR-expressing animals without PA (Supplemental Figure 5B). These results suggest that the delivery of GLR-1 synaptic sites is negatively regulated by local mitoROS production.

**Figure 4:**
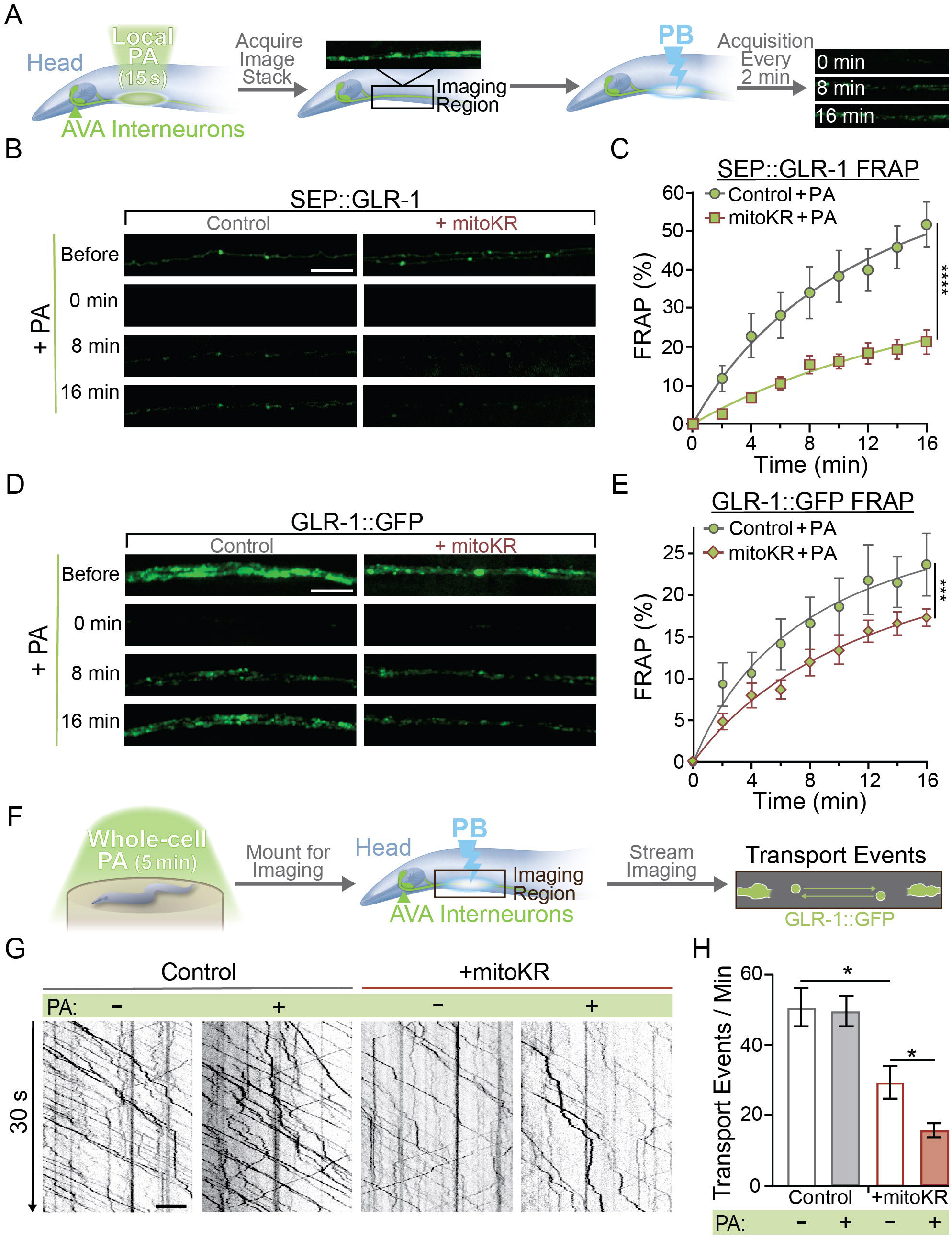
MitoROS downregulates the recruitment of GLR-1 to synapses. A) Diagram of experimental procedure followed for B-D (see Methods). B and D) Representative images of SEP::GLR-1 (B) or GLR-1::GFP (D) fluorescence before, immediately after, 8-, and 16-minutes after local PA and PB. C and E) Percent SEP (C) or GFP (E) fluorescence recovery after PB (FRAP) over 16 minutes after local PA and PB (n≥7 animals per group). ***: p<0.0005, ****: p<0.0001 using an extra sum-of-squares F-test with a Bonferroni correction. F) Diagram of experimental procedure followed for G-H (see Methods). G) 30-second representative kymographs of GLR-1::GFP movement in the AVA with or without global PA. H) Total number of transport events per minute quantified from 50-second-long kymographs (n=8 animals per group). All scale bars = 5 µm. Data is represented as mean ± s.e.m.; *: p<0.05 compared to controls or indicated experimental group using a One-Way ANOVA.

We speculated that ROS production by mitoKR could also impact transport of GLR-1 in a similar fashion to global ROS elevations shown previously^32^. Since local PA of mitoKR had no effect on the amount of GLR-1 transport (data not shown), we optimized a protocol to increase ROS production at mitochondria throughout AVA neurons. Whole-cell PA of mitoKR was achieved by illuminating freely behaving worms for 5 or 10 minutes. We used mito-roGFP to measure the resultant ROS increase at mitochondria from whole-cell PA and observed a slight increase in the average mito-roGFP F_ratio_ after 5 minutes of whole-cell PA and a significant increase in the F_ratio_ following a 10-minute whole-cell PA (Supplemental Figure 4E and 4F). Although not significantly increased from the unstimulated control, the 5-minute PA increased the F_ratio_ of mito-roGFP to 0.4, which is similar to the mito-roGFP F_ratio_ following 5 minutes of repetitive optical activation (Supplemental Figure 4F; Figure 3C). Therefore, we chose to do subsequent experiments using a whole-cell PA duration of 5 minutes. This whole-cell activation of mitoKR in the AVA prior to imaging GLR-1 transport (Figure 4F) revealed that cell-wide PA of mitoKR reduces the number of transport events (Figure 4G and 4H). These results coincide with our previous work showing that global elevations in ROS decrease export of GLR-1 out of the cell body^32^ and suggest that the mitochondria are a major source of the ROS involved in this regulation.

In summary, we have demonstrated that dendritic mitochondria take up Ca^2+^ in response to neuronal activity leading to an upregulation in ROS production at mitochondria. We also show that ROS production at mitochondria and loss or inhibition of MCU had opposite effects on GLR-1 exocytosis (Figure 2C and Figure 4C), so we hypothesized that local Ca^2+^ uptake by mitochondria and mitoROS production regulate the amount of GLR-1 exocytosis through the same signaling pathway. To test this, we subjected control or mitoKR-expressing worms to an acute Ru360 treatment, mounted them for imaging, and photoactivated a region of the AVA neurites prior to carrying out the FRAP protocol for SEP::GLR-1 (Figure 5A). This technique allowed us to bypass mitochondrial Ca^2+^ uptake and artificially induce ROS production at mitochondria in order to test if mitoROS is sufficient to downregulate synaptic recruitment of GLR-1. As shown previously, Ru360 treatment increased SEP::GLR-1 FRAP whereas mitoKR with local PA decreased the recovery rate. When Ru360 treatment was combined with local PA of mitoKR, the FRAP of SEP::GLR-1 was slightly delayed, but the relative fluorescence recovery after 16 minutes post-photobleach was nearly identical to local PA of mitoKR alone (Figure 5B and 5C). Interestingly, Ru360 treatment of mitoKR-expressing worms without PA had a FRAP rate that was comparable to the non-activated, untreated mitoKR group (Supplemental Figure 6A). Since artificial mitoROS production was able to occlude the effect of Ru360 on GLR-1 exocytosis, these results support that mitoROS is necessary and sufficient for downregulating GLR-1 exocytosis. Contrary to GLR-1 exocytosis, somatic export of GLR-1 is paradoxically reduced by both artificial mitoROS production and inhibition of MCU-1. To test if mitoROS and mitochondrial Ca^2+^ uptake regulate GLR-1 transport out of the cell body via the same mechanism, we combined acute Ru360 treatment with 5 minutes of whole-cell PA of mitoKR prior to imaging GLR-1 transport (Figure 5D). Both acute Ru360 treatment and whole-cell PA of mitoKR decreased the number of GLR-1 transport events to a similar extent (Figure 2G and 4H). When combined, the amount of GLR-1 transport was significantly decreased compared to Ru360 treatment alone and modestly decreased compared to mitoKR activation (Figure 5E and 5F). Ru360 treatment of mitoKR-expressing worms in the absence of PA had no additional effect on the amount of GLR-1 transport compared to untreated mitoKR-expressing worms (Supplemental Figure 6B). The compounding effect of mitoROS production and decreased mitochondrial Ca^2+^ uptake indicates that mitoROS signaling and mitochondrial Ca^2+^ uptake modulate GLR-1 transport via parallel regulatory pathways. This contrasts our observations of a Ca^2+^-dependent mitoROS signaling mechanism in the regulation of GLR-1 recruitment to synapses (Figure 5D) and suggests that mitochondrial activation and signaling varies based on subcellular location. Taken altogether, our results reveal a physiological mitoROS signaling mechanism that is initiated by activity-dependent Ca^2+^ uptake and downregulates GLR-1 recruitment to synapses.

**Figure 5:**
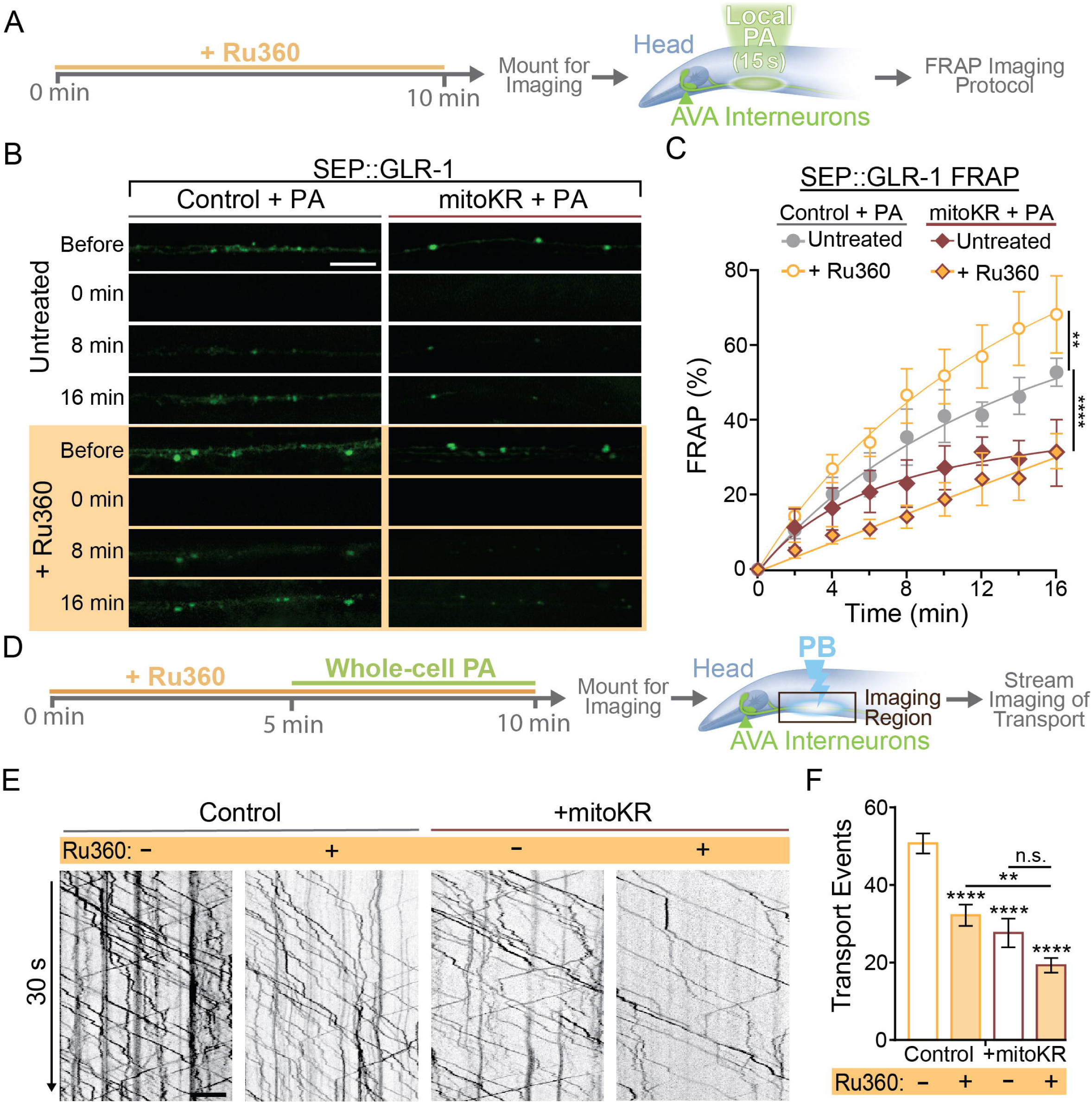
Regulation of synaptic recruitment of GLR-1 by mitoROS requires Ca^2+^ uptake via MCU-1. A) Diagram of experimental procedure in B-C (see Methods). B) Representative images of SEP fluorescence prior to, immediately after, and at 8- and 16-minutes post PB. C) Percent SEP FRAP over the 16-minute post PB (n≥5 animals per group). **: p<0.005, ****: p<0.0001 using an extra sum-of-squares F-test with a Bonferroni correction. D) Diagram of experimental procedure for E-F (see Methods). E) 30-second-long representative kymographs of GLR-1 transport for all + PA groups. F) Total number of transport events quantified from 50-second-long kymographs (n≥10 animals per group). All scale bars = 5 µm. Data is represented as mean ± s.e.m.; n.s = not significant, **: p<0.005, ****: p<0.0001 compared to controls or indicated experimental group using a One-Way ANOVA.

## DISCUSSION

We have outlined a novel activity-dependent mitochondrial signaling mechanism that negatively regulates excitatory synapse function. Our results support a model (Figure 6) in which synaptic activation leads to mitochondrial Ca^2+^ uptake via MCU-1 causing an increase in mitochondrial ROS that act to downregulate synaptic recruitment of AMPARs. The effect of mitoROS signaling on AMPAR recruitment to synapses appears to be due to the compounding effect of decreased transport out of the cell body, synaptic delivery as well as exocytosis of AMPARs to the synaptic membrane (Figure 4). This negative regulation by mitoROS may be a homeostatic mechanism that is important for prevention of excessive synaptic strengthening and the excitotoxicity that could result without this regulatory mechanism.

**Figure 6:**
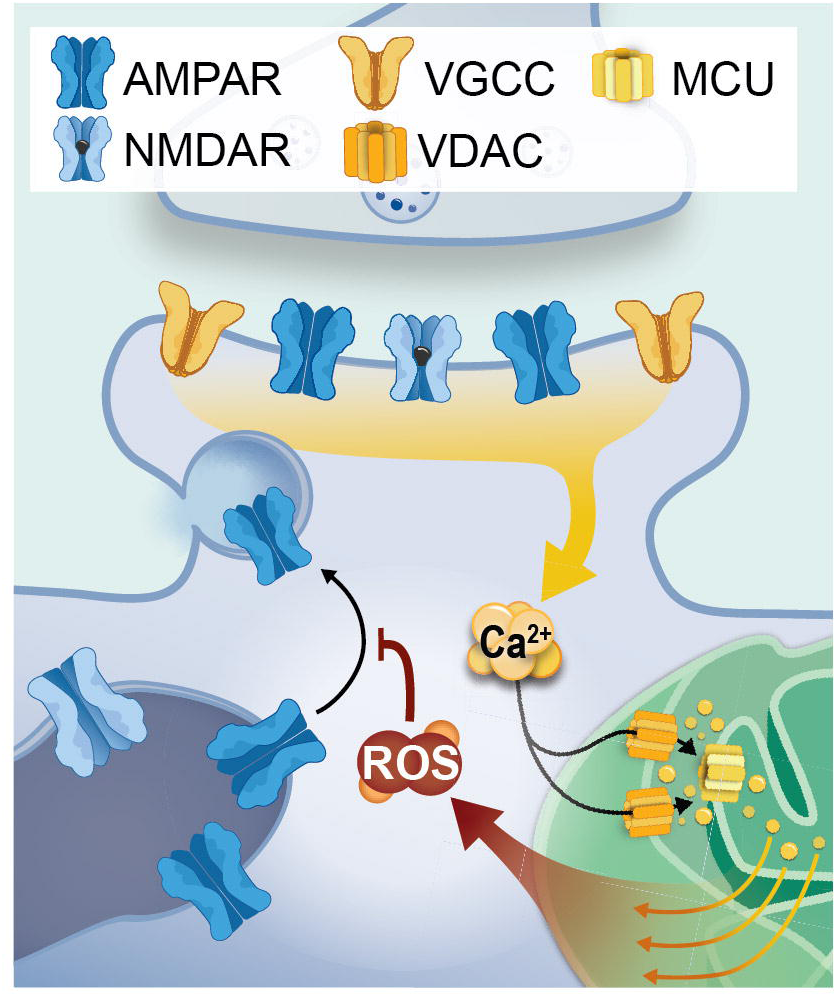
Model. In neurons, increased cytoplasmic Ca^2+^ due to activity-dependent opening of AMPARs, NMDARs and voltage-gated calcium channels (VGCCs) results in mitochondrial Ca^2+^ uptake via voltage-dependent anion channels (VDACs) at the outer mitochondrial membrane and further entry into the mitochondrial matrix via MCU. Once in the matrix, Ca^2+^ can directly and indirectly upregulate mitochondrial respiration from which ROS is a byproduct. The increased ROS can escape into the cytoplasm in the form of H_2_O_2_ and contribute to ROS signaling. Our data suggest that effectors of ROS signaling may be proteins involved in the exocytosis of AMPARs from transport vesicles or intracellular reserves (i.e., synaptic endosomes, left organelle) to the synaptic membrane.

### Mitochondrial calcium handling in synaptic function and plasticity

Buffering of cytoplasmic Ca^2+^ by mitochondria is thought to shape the spatiotemporal dynamics of Ca^2+^ signaling and upregulate mitochondrial output to meet energy demands^61^. Fine regulation of synaptic Ca^2+^ is particularly important because synaptic function and plasticity rely on a multitude of Ca^2+^-dependent signaling pathways that are all sensitive to the amplitude and duration of elevated Ca^2+^ ^15^. It is known that Ca^2+^ handling of presynaptic mitochondria modulates various presynaptic mechanisms central to synaptic transmission and plasticity including synaptic vesicle recycling^54, 56^ and release probability^49, 55, 62, 63^. Electron microscopy has revealed that mitochondria in the pre- and postsynaptic compartments of excitatory synapses differ in both size and electron density^34^ hinting that postsynaptic mitochondrial specialization is different than their presynaptic counterparts. However, only a few recently published studies have investigated if and how mitochondrial Ca^2+^ handling in dendrites regulates synaptic function or plasticity^64, 65^, and none have assessed the direct link between mitochondrial signaling and postsynaptic function in healthy neurons.

Postsynaptic plasticity mechanisms are also highly sensitive to the concentration and duration of elevated Ca^2+^ ^66, 67^, so Ca^2+^ uptake by postsynaptic mitochondria could shape Ca^2+^ events, and therefore synaptic transmission^65^. The importance of postsynaptic mitochondria for synaptic function could also be inferred from the decreased presence of synaptic mitochondria in Alzheimer’s and Parkinson’s disease that is observed before synaptic dysfunction^68^. Interestingly, mitochondrial transport in neurites is regulated by relative Ca^2+^ levels such that mitochondria deposition occurs at regions of high Ca^2+^, such as at pre- and postsynaptic sites^68^. If mitochondrial Ca^2+^ buffering truly contributes to cytoplasmic Ca^2+^ signaling, then one would expect an increase in cytoplasmic Ca^2+^ levels when mitochondrial Ca^2+^ uptake is diminished. It has been shown that loss of MCU-1 increases the amplitude and/or duration of cytoplasmic Ca^2+^ events in both invertebrate and vertebrate neurons ^64, 69, 70^. However, we did not detect a change in activity-dependent Ca^2+^ influx in the AVA neuron’s cytoplasm due to loss or inhibition of MCU-1 (Supplemental Figure 2F-2H). This discrepancy may be due to GCaMP6f’s high affinity for Ca^2+^ occluding slight changes in cytoplasmic Ca^2+^. Alternatively, mitochondrial Ca^2+^ uptake in AVA neurons, and perhaps *C. elegans* neurons in general, may be less reliant on MCU-1 function. It is also important to note that in our hands, the loss or pharmacological inhibition of MCU-1 did not completely abolish mitochondrial Ca^2+^ uptake. However, our observations are consistent with previous studies in which MCU-1 was conditionally or completely knocked out^51, 71^.

In addition to the importance of mitochondrial Ca^2+^ buffering for cytoplasmic signaling, there are many Ca^2+^-dependent processes within mitochondria. First, mitochondrial Ca^2+^ uptake can upregulate OXPHOS, and therefore ATP production^72^. It is possible that loss or inhibition of MCU-1 prevents activity-dependent upregulation of ATP which may indirectly impact endergonic mechanisms including GLR-1 transport, delivery and exocytosis^73–75^. However, our observations of upregulated GLR-1 delivery and exocytosis when MCU-1 is mutated or inhibited (Figure 2C) suggest that when mitochondrial Ca^2+^ uptake is decreased, ATP levels remain sufficient for local GLR-1 trafficking. Secondly, since ROS are a by-product of OXPHOS, ROS production can also be upregulated via several Ca^2+^-dependent mechanisms^76^. In fact, activity-induced mitoROS production via an MCU-1-dependent mechanism has been described in *C. elegans* in epidermal wound healing^38^. There is also evidence from *in vitro* studies in various human cell lines that MCU-dependent mitoROS signaling occurs in pathophysiological contexts such as during inflammation or hypoxia^77^. Lastly, mitochondrial Ca^2+^ uptake appears to be central to the pathophysiological plasticity mechanism that underlies hyperalgesia^78^. However, this work in addition to these previous studies prompt more questions than they answer in regard to postsynaptic roles of Ca^2+^-dependent mitoROS production.

### Regulation of AMPAR Trafficking by Mitochondrial ROS Signaling

The characteristics of ROS production and methods of action make them a diverse messenger molecule in various cell types, especially in the brain where metabolic activity and antioxidant mechanisms are higher than in other tissues^25, 79^. ROS signaling is compartmentalized due to the localization of ROS sources such as at the plasma membrane via NADPH oxidase or at mitochondria that is balanced by rapid cytoplasmic ROS scavenging^80, 81^. This is estimated to limit ROS diffusion to around 1 µm from its source^82^. Reversible protein oxidation by ROS is reminiscent of phosphorylation in that it can regulate protein folding, activation, and interactions^83^. Interestingly, the proportion of oxidizable protein residues is increased 4-fold in mammals compared to prokaryotes suggesting that ROS signaling may contribute to organismal complexity^84^.

Although mitochondria are regarded as the predominant source of ROS, there has been very little investigation of physiological mitoROS signaling in neurons *in vivo*. Recently, however, mitoROS production was shown to promote secretion of a neuropeptide from sensory neurons in *C. elegans* which activates antioxidant mechanisms in distal tissues^85^. There are also a few studies that demonstrate the functional relevance and versatility of mitoROS signaling in vertebrate neurons and their circuitry^86, 87^. Our results support an important mitochondrial signaling role and outline a mechanism in which activity-dependent mitoROS production regulates AMPAR localization. A comprehensive understanding of this mechanism would require systematically analyzing how protein oxidation alters the functionality of key players that regulate AMPAR delivery and recruitment to synapses.

There are several oxidizable candidate proteins and signaling molecules that regulate synaptic recruitment of AMPARs in neurons. Two major components of the Ca^2+^-signaling cascade that positively regulate AMPAR transport are calmodulin (CaM) and Ca^2+^/CaM-dependent protein kinase II (CaMKII) which are functionally regulated by oxidation^8, 11, 32^. CaM has two conserved methionines and when oxidized, the binding and activation of CaM to CaMKII is reduced^88^. When CaMKII is in its active Ca^2+^/CaM-bound conformation, oxidation of the regulatory domain enhances kinase activity^89^. Alternatively, when CaMKII is inactive, oxidation within the CaM binding domain prevents association of Ca^2+^/CaM with CaMKII^90^. At postsynaptic sites, recycling of AMPARs is regulated in a CaM/CaMKII-dependent manner meaning redox modification of these proteins can also influence AMPAR exocytosis and endocytosis at synapses^91^. Other proteins that regulate this process including protein kinase C^92^ (PKC) and the PDZ domain-containing scaffold protein Protein Interacting with C kinase 1^93^ (PICK-1). Activation of PKC following synaptic activation increases AMPAR insertion at synaptic membranes ^94^ whereas PICK-1 regulates AMPAR endocytosis^93^. Interestingly, ROS signaling can bi-directionally modulate PKC activity^95^ and oxidation of PICK-1 prevents its association with the synaptic membrane^96^. Although the effect of PICK-1 oxidation on synaptic expression of AMPARs has not been characterized, there is evidence that this redox mechanism regulates glutamatergic transmission and is protective during oxidative stress^97^. Thus, this current study opens the door to other questions regarding redox regulation of synaptic function and plasticity.

In contrast to the regulation of AMPAR exocytosis by postsynaptic mitoROS signaling (Figure 5B and 5C), we observed a compounding effect of MCU-1 inhibition and artificial mitoROS production on AMPAR export from the cell body (Figure 5E and 5F). These results suggest that AMPAR transport out of the cell body is regulated by mitochondrial Ca^2+^ handling and mitoROS production via two parallel signaling pathways. Since somatic mitochondria are morphologically distinct from their dendritic and axonal counterparts^98^, it is possible that they are functionally different as well. Altogether, these results open the door to questions regarding how functional diversity among mitochondria may allow mitochondrial signaling to differentially regulate signaling pathways based on subcellular location.

### Implications and Conclusion

Synaptic diversity is thought to enhance the computing power of the nervous system allowing for complex behaviors, a broad range of emotional states, and nearly endless memory storage. Interestingly, the proteomes of synaptic and non-synaptic mitochondria suggest that synaptic diversity may be enhanced by their resident mitochondria^99,100^. The proteomes of synaptic mitochondria allow for specialized function including activity-dependent regulation of ATP production and discrete Ca^2+^ handling abilities^18, 101^. The functional significance of enhanced energy capability and Ca^2+^ handling has been assessed for presynaptic mitochondria, but not in the context of postsynaptic sites. Here, we provide data indicating that postsynaptic mitochondria are functionally diverse and play a novel signaling role in regulating postsynaptic function.

In conclusion, we present evidence for a novel role for mitochondria in regulating the number of AMPARs at the synaptic membrane. This study proposes a mechanism in which Ca^2+^ signaling regulates mitochondrial ROS production providing a means of negative regulation of synaptic excitability in a way that may be important for synaptic homeostasis and prevention of excitotoxicity. This role for ROS signaling challenges the long-held misconception that elevated ROS is only detrimental to cells causing dysfunction and death^22^. Instead, mitoROS signaling acts as a physiological signal integrating synaptic function and mitochondrial output to link neuronal connectivity and metabolic capacity.

## EXPERIMENTAL PROCEDURES

### Plasmid Construction

See table S2 for details on plasmids used for this study. Plasmids were created using In-Fusion Cloning (Takara Bio) or the Gateway recombination (Invitrogen) method. DNA primers were created using Takara Bio’s online In-Fusion Primer Design Tool for In-Fusion Cloning and with the opensource ApE Plasmid Editor (M. Wayne Davis) for the Gateway recombination method.

### C. elegans Strains

*C. elegans* strains were maintained under standard conditions^102^(NGM with OP50 20°C). All animals used in experiments were day-one-old adult hermaphrodites which were selected 24 hours prior to experiments at the L4 stage. Transgenic strains (Table S1) were created by microinjection^103^ of *lin-15(n765ts)* worms with DNA mixes composed of the plasmids described in Table S2. All DNA mixes included a plasmid containing *lin-15(+)* to allow for phenotypic rescue of transgenic strains^104^. All strains used in optogenetic experiments were also mutant for the *lite-1* gene (allele: *ok530*) to limit off-target effects of our optical stimulation protocols due LITE-1^105^.

### Confocal Microscopy

All imaging was done using a Yokogawa CSUX1 spinning disc incorporated into a confocal microscope (Olympus IX83) with 405, 488, and 561 nm diode lasers (100-150 mW each; Andor ILE Laser Combiner). Images were captured using an Andor iXon Ultra EMCCD (DU-867) camera and a 100x/1.40NA oil objective (Olympus). Devices were controlled remotely for image acquisition using MetaMorph 7.10.1 (Molecular Devices).

### *In Vivo* Imaging of the AVA Neurites

One-day-old adult hermaphrodites were mounted for imaging by placing a single worm on an agar pad (10% agarose dissolved in M9 buffer) on a microscope slide with 1.6 µL of a solution containing equal measures of polystyrene beads (Polybead CAT No. 00876-15, Polysciences Inc.) and 30 mM muscimol (CAT No. 195336, MP Biomedicals). Once the muscimol slowed worm movement (∼5 minutes), a coverslip was dropped onto the agar pad physically restraining the worm. The worm’s orientation was manually adjusted by sliding the coverslip to reorient the positioning of the AVA interneurons for imaging^106^.

### Whole-Cell Neuronal Stimulation with ChRimson

Worms at the L4s stage from strains expressing ChRimson were picked onto an NGM/OP50 plate coated with a 100 µM concentration of all-Trans Retinal (Sigma-Aldrich, Cat No.: R2500-25; diluted with M9 buffer). Worms were left overnight on Retinal plates before optical neuronal activation via an LED array (613 nm, CoolBase 7 LED module from LuxeonStar). ChRimson expression was verified in these strains behaviorally by testing light-induced reversals (data not shown). For ChRimson activation before mito-roGFP imaging, freely behaving one-day-old adults were placed onto a fresh NGM/OP50 plate 2 inches beneath a 613 nm LED array. LED intensity was adjusted at the beginning of each experiment to 40 µW/mm^2^ using a custom potentiometer in combination with a digital optical power console (ThorLabs, PM100C) and photodiode sensor (ThorLabs, S170C). The pattern generator pulsed the LED for 1 s every 30 s (33.3 mHz) for 5 to 60 min before worms were mounted for imaging.

### Localized ChRimson Activation

To activate ChRimson within discrete regions of the AVA neurons, the neurites were located using a 100x objective, the co-expressed fluorescent reagents (i.e. mito-roGFP, GCaMP or mitoGCaMP) and the 488 nm imaging laser. Briefly, a fluorescent image of the co-expressed reagent in a single z-plane was acquired and a region mask was created on the AVA neurites. Then, the green LED (with a 605+20 nm filter; Chroma) from an LED illumination system (CoolLED pE300ultra) illuminated the masked region via projection through a Mosaic II digital mirror device (DMD; Andor Mosaic 3) controlled remotely via MetaMorph. LED intensity was adjusted to a total output of 5 µW using a digital optical power console (ThorLabs, PM100C) and microscope slide thermal sensor (ThorLabs, S175C). During the acquisition of an image stream, the master shutter of the DMD was controlled using MetaMorph’s “Trigger Components” function to illuminate the masked region for 3 seconds every 30 seconds.

### Ratiometric Fluorescence Imaging and Analysis of mito-roGFP

Immediately after ChRimson or mechano-stimulation, worms were mounted for imaging in a 15 mM Muscimol solution. The AVA neurites containing roGFP^+^ mitochondria were located, and images were collected with a 500 ms exposure every 0.25 µm to capture a stack of images (5.25 µm) around the neurites. The 525 nm emission was imaged with 405 nm then 488 nm illumination at each Z-plane. The average roGFP 525 nm fluorescence from 405 or 488 nm excitation was measured at individual mitochondria using MetaMorph’s region measurement tool in a single Z-plane where the roGFP fluorescence due to 488 nm excitation was the highest. The average background fluorescence near each mitochondrion was also collected. The mitochondria region trace was copied to the fluorescence image collected with 405 nm excitation at the corresponding Z-plane, then roGFP and background fluorescence values were logged.

### Whole-Cell mitoKR Activation

Individual one-day-old adults of transgenic strains (*csfEx168, csf195* or *csfEx188*) containing pRD36 [*Pflp-18::TOMM20::KillerRed::let-858*] as determined by the absence of the multi-vulva phenotype were transferred onto a fresh NGM/OP50 culture plate and placed 2 inches below a 567 nm LED array (CoolBase 7 LED module from LuxeonStar). The light intensity was adjusted to 25 µW/mm^2^ with our potentiometer, digital optical power console and photodiode sensor (S130C). Worms were illuminated for 5 or 10 minutes before being immediately mounted for imaging.

### Local mitoKR Activation

For localized photoactivation of mitoKR (TOMM20::KillerRed), the AVA neurites were located using a 100x objective, the co-expressed fluorescent reagents (i.e. mito-roGFP, GLR-1::GFP or SEP::GLR-1) and the 488 nm imaging laser. An image of mitoKR fluorescence in a single Z-plane was briefly acquired using a 100 ms exposure time and 561 nm imaging laser. Using this image, a region mask was created around a small region (100-300 µm^2^) containing mito-KR^+^ mitochondria. The green LED (with a 590+20 nm filter; Chroma) from our LED illumination system illuminated the masked region via projection of the green light through our DMD controlled by MetaMorph. LED intensity was adjusted to a total output of 10 µW using a digital optical power console (ThorLabs, PM100C) and photodiode sensor (ThorLabs, S130C). By remotely opening the DMD master shuttler, the masked region was illuminated for 15 s.

### Pharmacological inhibition of MCU-1 with Ru360

Ru360 (Sigma-Aldrich CAT No. 557440) was reconstituted in water at a concentration of 2 mM then distributed into 15 µL aliquots (in light safe microcentrifuge tubes) and stored at 4℃. Immediately before treatment, an Ru360 aliquot was diluted to 100 µM with M9 buffer. Then, 2-3 animals were placed on an NGM plate with OP50 and 200 µL of 100 µM Ru360 solution was pipetted onto the OP50 lawn where the animals resided, completely covering the lawn. Treatment was applied for 10 minutes after which point, the animal was removed to use in the outlined imaging protocols. For long-term optogenetic experiments, animals were bathed in the Ru360 treatment for ∼10 minutes before the Ru360-containing media naturally absorbed into the NGM/OP50 plate. The animals remained on this plate while undergoing the optical activation protocol for 5-60 minutes (see above sections, Whole-Cell Neuronal Stimulation with ChRimson and Whole-Cell mitoKR Activation).

### Transport Imaging and Analysis

All transport imaging was conducted on strains containing *akIs141* in the *glr-1* null background (*ky176*). The AVA neurites were located using the 100x objective and a 488 nm excitation laser to visualize GFP fluorescence. A consistent Z-plane was held in focus for the entire imaging session using the continuous focus function of a Z drift compensator (Olympus, IX3-ZDC2) controlled remotely by MetaMorph. Then, a proximal section of the neurites were photobleached using a 3 W, 488 nm Coherent solid-state laser (Genesis MX MTM; 0.5 W output; 1 s pulse) directed to the region defined in MetaMorph using a Mosaic II digital mirror device (Andor Mosaic 3). 30 seconds after photobleaching, an image stream was collected with the 488 nm excitation laser and a 100 ms exposure time. MetaMorph’s Kymograph tool was used to generate kymographs as previously reported ^6^. Transport events were quantified by manually counting all transport events from resultant kymographs.

### Fluorescence Recovery After Photobleaching (FRAP)

Strains expressing either GLR-1::GFP or SEP::GLR-1 were mounted for imaging as described above. Using the SEP or GFP fluorescence, a proximal region of the AVA neurites was localized. The stage position was memorized using MetaMorph’s stage position memory function and the ideal Z-plane was set using the ZDC control dialogue. An image stack SEP/GFP fluorescence was then acquired using the 488 nm excitation laser set to a 500 ms exposure. The Z-stack captures the entire width of the AVA process (21 z-planes; 0.25 µm steps, +/-2.5 µm around the neurite). If photoactivation was required for experiment, the shutter for the CoolLED system (pE-300^ultra^) was opened for the appropriate duration. Then, ∼40 µm sections of the neurite proximal and distal to the imaging region were photobleached using the same photobleaching settings as described for GLR-1 transport imaging. Lastly, the imaging region (40-50 µm) was photobleached. Immediately following, an image stack of SEP/GFP fluorescence was acquired for the 0-minute time point. Subsequent image stacks were acquired every 2 minutes out to 16 minutes. Resultant image stacks were processed and analyzed as previously described^32^ with the exception of the SEP FRAP dataset in Figure 2C. The individual timepoints in this dataset were not normalized to the initial fluorescence value per animal because initial SEP::GLR-1 fluorescence was significantly higher in *mcu-1(lf)* (Supplemental Figure 2B). Instead, the 0-minute fluorescence values were subtracted from the raw fluorescence values for all subsequent timepoints. Analysis of the fluorescence before photobleaching was analyzed by creating a data file (.log) of fluorescence values along the bleached region of the AVA neurite using MetaMorph’s linescan tool (line width=20 pixels). The resultant output file was analyzed using a custom MATLAB (R2021a) script to obtain the average area of fluorescent puncta (area under the peak).

### Imaging of mitoGCaMP

The AVA neurite was located and continuous autofocus was set as described above. Image streams (100 ms exposure) were collected with a 488 nm imaging laser (power=0.1%; attenuation=10). Localized ChRimson activation (see above section, Localized ChRimson Activation) was triggered every 30 seconds using MetaMorph’s “Trigger Components” feature starting 30 seconds after the start of the image stream. Imaging of mitoGCaMP fluorescence was continuous throughout the entire protocol covering all aspects of activation and rest.

### Experimental Design and Statistical Analyses

All relevant controls were included for each experimental replicate and all dataset were collected over 2-4 replicates. When manual quantification was required (i.e., for quantification of transport events from kymographs), the dataset was blinded to the genotype and experimental condition. Data was screened for outliers using the ROUT method (Q=1%). For FRAP datasets, animals were excluded if 50% or more of the timepoints were considered outliers. Experimental groups were considered significantly different if their comparison using a Student’s t-test (for comparing two groups) or One-way ANOVA with correction for multiple comparisons (Dunnett’s; for comparisons >2) yielded a p-value less than 0.05. To compare the FRAP rate between conditions, we used an extra sum-of-squares F-test comparing the best fit curve for each experimental group with a Bonferroni correction for multiple comparisons. Curves were considered different if a comparison yielded a p-value less than 0.01.

### Image and Data Presentation

All images were acquired under non-saturating conditions. Representative images were selected as they represent the average. Post-processing was done following analysis as needed to visualize corresponding quantifications. Image processing was performed in Photoshop (2023) and all images in each panel were identically processed. Graphs were created in GraphPad Prism (9.3.1) and exported as an enhanced metafile for integration into figures which were compiled in Adobe Illustrator (24.3). All data is represented as the mean ± the standard error of the mean. Illustrations were created in their entirety in Adobe Illustrator.

### Code/Software

Custom Excel modules (created in Excel’s Visual Basic Editor) were used for analysis of cytoplasmic and mitochondrial calcium imaging. The modules are available online at: https://github.com/racheldoser/GCaMP_Analysis_Excel_VBA.git.

## ACKNOWLEDGEMENTS

This work was supported in part by an R01 from NIH/NINDS(NS159), we thank Sasha de Henau for the mito-roGFP plasmid; Attila Stetak for the GCaMP6f plasmid and the CGC at UMN for strains.

## AUTHOR CONTRIBUTIONS

R.D., K.K. and F.H. designed the experiments and wrote the manuscript. E.D. compiled the supplemental data. R.D., K.K. and E.D. created the genetic reagents, transgenic strains and analyzed data. R.D. created the figures and illustrations. R.D., F.H., K.K., and E.D. edited the manuscript.

## REFERENCES

1. Niciu, M.J., Kelmendi, B., and Sanacora, G. (2012). Overview of glutamatergic neurotransmission in the nervous system. Pharmacol Biochem Behav 100, 656– 664. 10.1016/j.pbb.2011.08.008.

2. Popoli, M., Yan, Z., McEwen, B.S., and Sanacora, G. (2012). The stressed synapse: the impact of stress and glucocorticoids on glutamate transmission. Nat Rev Neurosci 13, 22–37. 10.1038/nrn3138.

3. Schaeuble, D., Packard, A.E.B., McKlveen, J.M., Morano, R., Fourman, S., Smith, B.L., Scheimann, J.R., Packard, B.A., Wilson, S.P., James, J., et al. (2019). Prefrontal Cortex Regulates Chronic Stress-Induced Cardiovascular Susceptibility. J Am Heart Assoc 8. 10.1161/JAHA.119.014451.

4. Kim, C.-H., and Lisman, J.E. (2001). A Labile Component of AMPA Receptor-Mediated Synaptic Transmission Is Dependent on Microtubule Motors, Actin, and N -Ethylmaleimide-Sensitive Factor. The Journal of Neuroscience 21, 4188–4194. 10.1523/JNEUROSCI.21-12-04188.2001.

5. Setou, M., Seog, D.-H., Tanaka, Y., Kanai, Y., Takei, Y., Kawagishi, M., and Hirokawa, N. (2002). Glutamate-receptor-interacting protein GRIP1 directly steers kinesin to dendrites. Nature 417, 83–87. 10.1038/nature743.

6. Hoerndli, F.J., Maxfield, D.A., Brockie, P.J., Mellem, J.E., Jensen, E., Wang, R., Madsen, D.M., and Maricq, A. V. (2013). Kinesin-1 Regulates Synaptic Strength by Mediating the Delivery, Removal, and Redistribution of AMPA Receptors. Neuron 80, 1421–1437. 10.1016/j.neuron.2013.10.050.

7. Esteves da Silva, M., Adrian, M., Schätzle, P., Lipka, J., Watanabe, T., Cho, S., Futai, K., Wierenga, C.J., Kapitein, L.C., and Hoogenraad, C.C. (2015). Positioning of AMPA Receptor-Containing Endosomes Regulates Synapse Architecture. Cell Rep 13, 933–943. 10.1016/j.celrep.2015.09.062.

8. Hangen, E., Cordelières, F.P., Petersen, J.D., Choquet, D., and Coussen, F. (2018). Neuronal Activity and Intracellular Calcium Levels Regulate Intracellular Transport of Newly Synthesized AMPAR. Cell Rep 24, 1001–1012.e3. 10.1016/j.celrep.2018.06.095.

9. Hoerndli, F.J., Brockie, P.J., Wang, R., Mellem, J.E., Kallarackal, A., Doser, R.L., Pierce, D.M., Madsen, D.M., and Maricq, A. V. (2022). MAPK signaling and a mobile scaffold complex regulate AMPA receptor transport to modulate synaptic strength. Cell Rep 38, 110577. 10.1016/j.celrep.2022.110577.

10. Yang, Y., Wang, X., Frerking, M., and Zhou, Q. (2008). Delivery of AMPA receptors to perisynaptic sites precedes the full expression of long-term potentiation. Proceedings of the National Academy of Sciences 105, 11388– 11393. 10.1073/pnas.0802978105.

11. Hoerndli, F.J., Wang, R., Mellem, J.E., Kallarackal, A., Brockie, P.J., Thacker, C., Madsen, D.M., and Maricq, A. V. (2015). Neuronal Activity and CaMKII Regulate Kinesin-Mediated Transport of Synaptic AMPARs. Neuron 86, 457–474. 10.1016/j.neuron.2015.03.011.

12. Ehlers, M.D. (2000). Reinsertion or Degradation of AMPA Receptors Determined by Activity-Dependent Endocytic Sorting. Neuron 28, 511–525. 10.1016/S0896-6273(00)00129-X.

13. Yudowski, G.A., Puthenveedu, M.A., Leonoudakis, D., Panicker, S., Thorn, K.S., Beattie, E.C., and von Zastrow, M. (2007). Real-Time Imaging of Discrete Exocytic Events Mediating Surface Delivery of AMPA Receptors. Journal of Neuroscience 27, 11112–11121. 10.1523/JNEUROSCI.2465-07.2007.

14. Choquet, D., and Triller, A. (2013). The Dynamic Synapse. Neuron 80, 691–703. 10.1016/j.neuron.2013.10.013.

15. Nakahata, Y., and Yasuda, R. (2018). Plasticity of Spine Structure: Local Signaling, Translation and Cytoskeletal Reorganization. Front Synaptic Neurosci 10. 10.3389/fnsyn.2018.00029.

16. Gutiérrez, Y., López-García, S., Lario, A., Gutiérrez-Eisman, S., Delevoye, C., and Esteban, J.A. (2021). KIF13A drives AMPA receptor synaptic delivery for long-term potentiation via endosomal remodeling. Journal of Cell Biology 220. 10.1083/jcb.202003183.

17. Wacquier, B., Combettes, L., and Dupont, G. (2019). Cytoplasmic and Mitochondrial Calcium Signaling: A Two-Way Relationship. Cold Spring Harb Perspect Biol 11, a035139. 10.1101/cshperspect.a035139.

18. Faria-Pereira, A., and Morais, V.A. (2022). Synapses: The Brain’s Energy-Demanding Sites. Int J Mol Sci 23, 3627. 10.3390/ijms23073627.

19. Chae, S., Ahn, B.Y., Byun, K., Cho, Y.M., Yu, M.-H., Lee, B., Hwang, D., and Park, K.S. (2013). A Systems Approach for Decoding Mitochondrial Retrograde Signaling Pathways. Sci Signal 6. 10.1126/scisignal.2003266.

20. Hirabayashi, Y., Kwon, S.-K., Paek, H., Pernice, W.M., Paul, M.A., Lee, J., Erfani, P., Raczkowski, A., Petrey, D.S., Pon, L.A., et al. (2017). ER-mitochondria tethering by PDZD8 regulates Ca2+ dynamics in mammalian neurons. Science (1979) 358, 623–630. 10.1126/science.aan6009.

21. Angelova, P.R., and Abramov, A.Y. (2018). Role of mitochondrial ROS in the brain: from physiology to neurodegeneration. FEBS Lett 592, 692–702. 10.1002/1873-3468.12964.

22. Sies, H., and Jones, D.P. (2020). Reactive oxygen species (ROS) as pleiotropic physiological signalling agents. Nat Rev Mol Cell Biol 21, 363–383. 10.1038/s41580-020-0230-3.

23. Hidalgo, C., and Arias-Cavieres, A. (2016). Calcium, reactive oxygen species, and synaptic plasticity. Physiology 31, 201–215. 10.1152/physiol.00038.2015.

24. Oswald, M.C.W., Garnham, N., Sweeney, S.T., and Landgraf, M. (2018). Regulation of neuronal development and function by ROS. FEBS Lett 592, 679– 691. 10.1002/1873-3468.12972.

25. Biswas, K., Alexander, K., and Francis, M.M. (2022). Reactive Oxygen Species: Angels and Demons in the Life of a Neuron. NeuroSci 3, 130–145. 10.3390/neurosci3010011.

26. Massaad, C.A., and Klann, E. (2011). Reactive oxygen species in the regulation of synaptic plasticity and memory. Antioxid Redox Signal 14, 2013–2054. 10.1089/ars.2010.3208.

27. Oswald, M.C.W., Brooks, P.S., Zwart, M.F., Mukherjee, A., West, R.J.H., Giachello, C.N.G., Morarach, K., Baines, R.A., Sweeney, S.T., and Landgraf, M. (2018). Reactive oxygen species regulate activity-dependent neuronal plasticity in Drosophila. Elife 7. 10.7554/eLife.39393.

28. Klann, E., Roberson, E.D., Knapp, L.T., and Sweatt, J.D. (1998). A role for superoxide in protein kinase C activation and induction of long-term potentiation. J Biol Chem 273, 4516–4522. 10.1074/jbc.273.8.4516.

29. Knapp, L.T., and Klann, E. (2002). Potentiation of Hippocampal Synaptic Transmission by Superoxide Requires the Oxidative Activation of Protein Kinase C. The Journal of Neuroscience 22, 674–683. 10.1523/JNEUROSCI.22-03-00674.2002.

30. Huddleston, A.T., Tang, W., Takeshima, H., Hamilton, S.L., and Klann, E. (2008). Superoxide-Induced Potentiation in the Hippocampus Requires Activation of Ryanodine Receptor Type 3 and ERK. J Neurophysiol 99, 1565–1571. 10.1152/jn.00659.2007.

31. Lee, D.Z., Chung, J.M., Chung, K., and Kang, M.-G. (2012). Reactive oxygen species (ROS) modulate AMPA receptor phosphorylation and cell-surface localization in concert with pain-related behavior. Pain 153, 1905–1915. 10.1016/j.pain.2012.06.001.

32. Doser, R.L., Amberg, G.C., and Hoerndli, F.J. (2020). Reactive Oxygen Species Modulate Activity-Dependent AMPA Receptor Transport in C. elegans. The Journal of Neuroscience 40, 7405–7420. 10.1523/JNEUROSCI.0902-20.2020.

33. Doser, R.L., and Hoerndli, F.J. (2022). Decreased Reactive Oxygen Species Signaling Alters Glutamate Receptor Transport to Synapses in C. elegans AVA Neurons. MicroPubl Biol 2022. 10.17912/micropub.biology.000528.

34. Freeman, D., Petralia, R., Wang, Y.-X., Mattson, M., and Yao, P. (2017). Mitochondria in hippocampal presynaptic and postsynaptic compartments differ in size as well as intensity. Matters (Zur). 10.19185/matters.201711000009.

35. Back, P., Braeckman, B.P., and Matthijssens, F. (2012). ROS in Aging Caenorhabditis elegansL: Damage or Signaling? Oxid Med Cell Longev 2012, 1–14. 10.1155/2012/608478.

36. Petriv, O.I., and Rachubinski, R.A. (2004). Lack of Peroxisomal Catalase Causes a Progeric Phenotype in Caenorhabditis elegans. Journal of Biological Chemistry 279, 19996–20001. 10.1074/jbc.M400207200.

37. Morsci, N.S., Hall, D.H., Driscoll, M., and Sheng, Z.-H. (2016). Age-Related Phasic Patterns of Mitochondrial Maintenance in Adult *Caenorhabditis elegans* Neurons. The Journal of Neuroscience 36, 1373–1385. 10.1523/JNEUROSCI.2799-15.2016.

38. Xu, S., and Chisholm, A.D. (2014). C. elegans Epidermal Wounding Induces a Mitochondrial ROS Burst that Promotes Wound Repair. Dev Cell 31, 48–60. 10.1016/j.devcel.2014.08.002.

39. Alvarez, J., Alvarez-Illera, P., García-Casas, P., Fonteriz, R.I., and Montero, M. (2020). The Role of Ca2+ Signaling in Aging and Neurodegeneration: Insights from Caenorhabditis elegans Models. Cells 9, 204. 10.3390/cells9010204.

40. Sengupta, P., and Samuel, A.D. (2009). Caenorhabditis elegans: a model system for systems neuroscience. Curr Opin Neurobiol 19, 637–643. 10.1016/j.conb.2009.09.009.

41. Cook, S.J., Jarrell, T.A., Brittin, C.A., Wang, Y., Bloniarz, A.E., Yakovlev, M.A., Nguyen, K.C.Q., Tang, L.T.-H., Bayer, E.A., Duerr, J.S., et al. (2019). Whole-animal connectomes of both Caenorhabditis elegans sexes. Nature 571, 63–71. 10.1038/s41586-019-1352-7.

42. Taylor, S.R., Santpere, G., Weinreb, A., Barrett, A., Reilly, M.B., Xu, C., Varol, E., Oikonomou, P., Glenwinkel, L., McWhirter, R., et al. (2021). Molecular topography of an entire nervous system. Cell 184, 4329–4347.e23. 10.1016/j.cell.2021.06.023.

43. Maricq, A. V., Peckol, E., Driscoll, M., and Bargmann, C.I. (1995). Mechanosensory signalling in C. elegans mediated by the GLR-1 glutamate receptor. Nature 378, 78–81. 10.1038/378078a0.

44. Rongo, C., and Kaplan, J.M. (1999). CaMKII regulates the density of central glutamatergic synapses in vivo. Nature 402, 195–199. 10.1038/46065.

45. Widagdo, J., Guntupalli, S., Jang, S.E., and Anggono, V. (2017). Regulation of AMPA Receptor Trafficking by Protein Ubiquitination. Front Mol Neurosci 10. 10.3389/fnmol.2017.00347.

46. Brini, M., Calì, T., Ottolini, D., and Carafoli, E. (2013). Intracellular Calcium Homeostasis and Signaling. In, pp. 119–168. 10.1007/978-94-007-5561-1_5.

47. Márkus, N.M., Hasel, P., Qiu, J., Bell, K.F.S., Heron, S., Kind, P.C., Dando, O., Simpson, T.I., and Hardingham, G.E. (2016). Expression of mRNA Encoding Mcu and Other Mitochondrial Calcium Regulatory Genes Depends on Cell Type, Neuronal Subtype, and Ca2+ Signaling. PLoS One 11, e0148164. 10.1371/journal.pone.0148164.

48. Klapoetke, N.C., Murata, Y., Kim, S.S., Pulver, S.R., Birdsey-Benson, A., Cho, Y.K., Morimoto, T.K., Chuong, A.S., Carpenter, E.J., Tian, Z., et al. (2014). Independent optical excitation of distinct neural populations. Nat Methods 11, 338–346. 10.1038/nmeth.2836.

49. Ashrafi, G., de Juan-Sanz, J., Farrell, R.J., and Ryan, T.A. (2020). Molecular Tuning of the Axonal Mitochondrial Ca2+ Uniporter Ensures Metabolic Flexibility of Neurotransmission. Neuron 105, 678–687.e5. 10.1016/j.neuron.2019.11.020.

50. Baughman, J.M., Perocchi, F., Girgis, H.S., Plovanich, M., Belcher-Timme, C.A., Sancak, Y., Bao, X.R., Strittmatter, L., Goldberger, O., Bogorad, R.L., et al. (2011). Integrative genomics identifies MCU as an essential component of the mitochondrial calcium uniporter. Nature 476, 341–345. 10.1038/nature10234.

51. Álvarez-Illera, P., García-Casas, P., Fonteriz, R.I., Montero, M., and Alvarez, J. (2020). Mitochondrial Ca2+ Dynamics in MCU Knockout C. elegans Worms. Int J Mol Sci 21, 8622. 10.3390/ijms21228622.

52. Woods, J.J., Nemani, N., Shanmughapriya, S., Kumar, A., Zhang, M., Nathan, S.R., Thomas, M., Carvalho, E., Ramachandran, K., Srikantan, S., et al. (2019). A Selective and Cell-Permeable Mitochondrial Calcium Uniporter (MCU) Inhibitor Preserves Mitochondrial Bioenergetics after Hypoxia/Reoxygenation Injury. ACS Cent Sci 5, 153–166. 10.1021/acscentsci.8b00773.

53. Stoler, O., Stavsky, A., Khrapunsky, Y., Melamed, I., Stutzmann, G., Gitler, D., Sekler, I., and Fleidervish, I. (2022). Frequency- and spike-timing-dependent mitochondrial Ca2+ signaling regulates the metabolic rate and synaptic efficacy in cortical neurons. Elife 11. 10.7554/eLife.74606.

54. Billups, B., and Forsythe, I.D. (2002). Presynaptic Mitochondrial Calcium Sequestration Influences Transmission at Mammalian Central Synapses. The Journal of Neuroscience 22, 5840–5847. 10.1523/JNEUROSCI.22-14-05840.2002.

55. Sun, T., Qiao, H., Pan, P.-Y., Chen, Y., and Sheng, Z.-H. (2013). Motile Axonal Mitochondria Contribute to the Variability of Presynaptic Strength. Cell Rep 4, 413–419. 10.1016/j.celrep.2013.06.040.

56. Marland, J.R.K., Hasel, P., Bonnycastle, K., and Cousin, M.A. (2016). Mitochondrial Calcium Uptake Modulates Synaptic Vesicle Endocytosis in Central Nerve Terminals. Journal of Biological Chemistry 291, 2080–2086. 10.1074/jbc.M115.686956.

57. Petrini, E.M., Lu, J., Cognet, L., Lounis, B., Ehlers, M.D., and Choquet, D. (2009). Endocytic Trafficking and Recycling Maintain a Pool of Mobile Surface AMPA Receptors Required for Synaptic Potentiation. Neuron 63, 92–105. 10.1016/j.neuron.2009.05.025.

58. Morgan, B., Sobotta, M.C., and Dick, T.P. (2011). Measuring EGSH and H2O2 with roGFP2-based redox probes. Free Radic Biol Med 51, 1943–1951. 10.1016/j.freeradbiomed.2011.08.035.

59. Schafer, W.R. (2015). Mechanosensory molecules and circuits in C. elegans. Pflugers Arch 467, 39–48. 10.1007/s00424-014-1574-3.

60. Braeckman, B.P., Smolders, A., Back, P., and De Henau, S. (2016). In Vivo Detection of Reactive Oxygen Species and Redox Status in Caenorhabditis elegans. Antioxid Redox Signal 25, 577–592. 10.1089/ars.2016.6751.

61. Duchen, M.R. (2000). Mitochondria and calcium: from cell signalling to cell death. J Physiol 529, 57–68. 10.1111/j.1469-7793.2000.00057.x.

62. Lee, D., Lee, K.-H., Ho, W.-K., and Lee, S.-H. (2007). Target Cell-Specific Involvement of Presynaptic Mitochondria in Post-Tetanic Potentiation at Hippocampal Mossy Fiber Synapses. The Journal of Neuroscience 27, 13603– 13613. 10.1523/JNEUROSCI.3985-07.2007.

63. Devine, M.J., Szulc, B.R., Howden, J.H., López-Doménech, G., Ruiz, A., and Kittler, J.T. (2022). Mitochondrial Ca2+ uniporter haploinsufficiency enhances long-term potentiation at hippocampal mossy fibre synapses. J Cell Sci 135. 10.1242/jcs.259823.

64. Groten, C.J., and MacVicar, B.A. (2022). Mitochondrial Ca2+ uptake by the MCU facilitates pyramidal neuron excitability and metabolism during action potential firing. Commun Biol 5, 900. 10.1038/s42003-022-03848-1.

65. O’Hare, J.K., Gonzalez, K.C., Herrlinger, S.A., Hirabayashi, Y., Hewitt, V.L., Blockus, H., Szoboszlay, M., Rolotti, S. V., Geiller, T.C., Negrean, A., et al. (2022). Compartment-specific tuning of dendritic feature selectivity by intracellular Ca ^2+^ release. Science (1979) 375. 10.1126/science.abm1670.

66. Huganir, R.L., and Nicoll, R.A. (2013). AMPARs and Synaptic Plasticity: The Last 25 Years. Neuron 80, 704–717. 10.1016/j.neuron.2013.10.025.

67. Citri, A., and Malenka, R.C. (2008). Synaptic Plasticity: Multiple Forms, Functions, and Mechanisms. Neuropsychopharmacology 33, 18–41. 10.1038/sj.npp.1301559.

68. Sheng, Z.-H. (2014). Mitochondrial trafficking and anchoring in neurons: New insight and implications. Journal of Cell Biology 204, 1087–1098. 10.1083/jcb.201312123.

69. Bisbach, C.M., Hutto, R.A., Poria, D., Cleghorn, W.M., Abbas, F., Vinberg, F., Kefalov, V.J., Hurley, J.B., and Brockerhoff, S.E. (2020). Mitochondrial Calcium Uniporter (MCU) deficiency reveals an alternate path for Ca2+ uptake in photoreceptor mitochondria. Sci Rep 10, 16041. 10.1038/s41598-020-72708-x.

70. Nichols, M., Elustondo, P.A., Warford, J., Thirumaran, A., Pavlov, E. V, and Robertson, G.S. (2017). Global ablation of the mitochondrial calcium uniporter increases glycolysis in cortical neurons subjected to energetic stressors. Journal of Cerebral Blood Flow & Metabolism 37, 3027–3041. 10.1177/0271678X16682250.

71. Hamilton, J., Brustovetsky, T., Rysted, J.E., Lin, Z., Usachev, Y.M., and Brustovetsky, N. (2018). Deletion of mitochondrial calcium uniporter incompletely inhibits calcium uptake and induction of the permeability transition pore in brain mitochondria. Journal of Biological Chemistry 293, 15652–15663. 10.1074/jbc.RA118.002926.

72. Rossi, A., Pizzo, P., and Filadi, R. (2019). Calcium, mitochondria and cell metabolism: A functional triangle in bioenergetics. Biochimica et Biophysica Acta (BBA) - Molecular Cell Research 1866, 1068–1078. 10.1016/j.bbamcr.2018.10.016.

73. Schnitzer, M.J., and Block, S.M. (1997). Kinesin hydrolyses one ATP per 8-nm step. Nature 388, 386–390. 10.1038/41111.

74. Hanley, J.G. (2007). NSF binds calcium to regulate its interaction with AMPA receptor subunit GluR2. J Neurochem 101, 1644–1650. 10.1111/j.1471-4159.2007.04455.x.

75. Araki, Y., Lin, D.-T., and Huganir, R.L. (2010). Plasma membrane insertion of the AMPA receptor GluA2 subunit is regulated by NSF binding and Q/R editing of the ion pore. Proceedings of the National Academy of Sciences 107, 11080–11085. 10.1073/pnas.1006584107.

76. Görlach, A., Bertram, K., Hudecova, S., and Krizanova, O. (2015). Calcium and ROS: A mutual interplay. Redox Biol 6, 260–271. 10.1016/j.redox.2015.08.010.

77. Dong, Z., Shanmughapriya, S., Tomar, D., Siddiqui, N., Lynch, S., Nemani, N., Breves, S.L., Zhang, X., Tripathi, A., Palaniappan, P., et al. (2017). Mitochondrial Ca2+ Uniporter Is a Mitochondrial Luminal Redox Sensor that Augments MCU Channel Activity. Mol Cell 65, 1014–1028.e7. 10.1016/j.molcel.2017.01.032.

78. Kim, H.Y., Lee, K.Y., Lu, Y., Wang, J., Cui, L., Kim, S.J., Chung, J.M., and Chung, K. (2011). Mitochondrial Ca ^2+^ Uptake Is Essential for Synaptic Plasticity in Pain. The Journal of Neuroscience 31, 12982–12991. 10.1523/JNEUROSCI.3093-11.2011.

79. Vicente-Gutiérrez, C., Jiménez-Blasco, D., and Quintana-Cabrera, R. (2021). Intertwined ROS and Metabolic Signaling at the Neuron-Astrocyte Interface. Neurochem Res 46, 23–33. 10.1007/s11064-020-02965-9.

80. Niemeyer, J., Scheuring, D., Oestreicher, J., Morgan, B., and Schroda, M. (2021). Real-time monitoring of subcellular H2O2 distribution in *Chlamydomonas reinhardtii*. Plant Cell 33, 2935–2949. 10.1093/plcell/koab176.

81. Sies, H. (2017). Hydrogen peroxide as a central redox signaling molecule in physiological oxidative stress: Oxidative eustress. Redox Biol 11, 613–619. 10.1016/j.redox.2016.12.035.

82. Lim, J.B., Huang, B.K., Deen, W.M., and Sikes, H.D. (2015). Analysis of the lifetime and spatial localization of hydrogen peroxide generated in the cytosol using a reduced kinetic model. Free Radic Biol Med 89, 47–53. 10.1016/j.freeradbiomed.2015.07.009.

83. Miseta, A., and Csutora, P. (2000). Relationship Between the Occurrence of Cysteine in Proteins and the Complexity of Organisms. Mol Biol Evol 17, 1232– 1239. 10.1093/oxfordjournals.molbev.a026406.

84. Go, Y.-M., and Jones, D.P. (2013). The Redox Proteome. Journal of Biological Chemistry 288, 26512–26520. 10.1074/jbc.R113.464131.

85. Jia, Q., and Sieburth, D. (2021). Mitochondrial hydrogen peroxide positively regulates neuropeptide secretion during diet-induced activation of the oxidative stress response. Nat Commun 12, 2304. 10.1038/s41467-021-22561-x.

86. Bao, L., Avshalumov, M. V., Patel, J.C., Lee, C.R., Miller, E.W., Chang, C.J., and Rice, M.E. (2009). Mitochondria Are the Source of Hydrogen Peroxide for Dynamic Brain-Cell Signaling. Journal of Neuroscience 29, 9002–9010. 10.1523/JNEUROSCI.1706-09.2009.

87. Accardi, M. V., Daniels, B.A., Brown, P.M.G.E., Fritschy, J.-M., Tyagarajan, S.K., and Bowie, D. (2014). Mitochondrial reactive oxygen species regulate the strength of inhibitory GABA-mediated synaptic transmission. Nat Commun 5, 3168. 10.1038/ncomms4168.

88. Robison, A.J., Winder, D.G., Colbran, R.J., and Bartlett, R.K. (2007). Oxidation of calmodulin alters activation and regulation of CaMKII. Biochem Biophys Res Commun 356, 97–101. 10.1016/j.bbrc.2007.02.087.

89. Erickson, J.R., Joiner, M. ling A., Guan, X., Kutschke, W., Yang, J., Oddis, C. V., Bartlett, R.K., Lowe, J.S., O’Donnell, S.E., Aykin-Burns, N., et al. (2008). A Dynamic Pathway for Calcium-Independent Activation of CaMKII by Methionine Oxidation. Cell 133, 462–474. 10.1016/j.cell.2008.02.048.

90. Konstantinidis, K., Bezzerides, V.J., Lai, L., Isbell, H.M., Wei, A.-C., Wu, Y., Viswanathan, M.C., Blum, I.D., Granger, J.M., Heims-Waldron, D., et al. (2020). MICAL1 constrains cardiac stress responses and protects against disease by oxidizing CaMKII. Journal of Clinical Investigation 130, 4663–4678. 10.1172/JCI133181.

91. Bayer, K.U., and Schulman, H. (2019). CaM Kinase: Still Inspiring at 40. Neuron 103, 380–394. 10.1016/j.neuron.2019.05.033.

92. Boehm, J., Kang, M.G., Johnson, R.C., Esteban, J., Huganir, R.L., and Malinow, R. (2006). Synaptic Incorporation of AMPA Receptors during LTP Is Controlled by a PKC Phosphorylation Site on GluR1. Neuron 51, 213–225. 10.1016/j.neuron.2006.06.013.

93. Fiuza, M., Rostosky, C.M., Parkinson, G.T., Bygrave, A.M., Halemani, N., Baptista, M., Milosevic, I., and Hanley, J.G. (2017). PICK1 regulates AMPA receptor endocytosis via direct interactions with AP2 α-appendage and dynamin. Journal of Cell Biology 216, 3323–3338. 10.1083/jcb.201701034.

94. Ren, S.Q., Yan, J.Z., Zhang, X.Y., Bu, Y.F., Pan, W.W., Yao, W., Tian, T., and Lu, W. (2013). PKCλ is critical in AMPA receptor phosphorylation and synaptic incorporation during LTP. EMBO Journal 32, 1365–1380. 10.1038/emboj.2013.60.

95. Steinberg, S.F. (2015). Mechanisms for redox-regulation of protein kinase C. Front Pharmacol 6. 10.3389/fphar.2015.00128.

96. Shi, Y., Yu, J., Jia, Y., Pan, L., Shen, C., Xia, J., and Zhang, M. (2010). Redox-Regulated Lipid Membrane Binding of the PICK1 PDZ Domain. Biochemistry 49, 4432–4439. 10.1021/bi100269t.

97. Wang, Y.-N., Zhou, L., Li, Y.-H., Wang, Z., Li, Y.-C., Zhang, Y.-W., Wang, Y., Liu, G., and Shen, Y. (2015). Protein Interacting with C-Kinase 1 Deficiency Impairs Glutathione Synthesis and Increases Oxidative Stress via Reduction of Surface Excitatory Amino Acid Carrier 1. The Journal of Neuroscience 35, 6429–6443. 10.1523/JNEUROSCI.3966-14.2015.

98. Lee, A., Hirabayashi, Y., Kwon, S.-K., Lewis, T.L., and Polleux, F. (2018). Emerging roles of mitochondria in synaptic transmission and neurodegeneration. Curr Opin Physiol 3, 82–93. 10.1016/j.cophys.2018.03.009.

99. Stauch, K.L., Purnell, P.R., and Fox, H.S. (2014). Quantitative Proteomics of Synaptic and Nonsynaptic Mitochondria: Insights for Synaptic Mitochondrial Vulnerability. J Proteome Res 13, 2620–2636. 10.1021/pr500295n.

100. Graham, L.C., Eaton, S.L., Brunton, P.J., Atrih, A., Smith, C., Lamont, D.J., Gillingwater, T.H., Pennetta, G., Skehel, P., and Wishart, T.M. (2017). Proteomic profiling of neuronal mitochondria reveals modulators of synaptic architecture. Mol Neurodegener 12, 77. 10.1186/s13024-017-0221-9.

101. Brown, M.R., Sullivan, P.G., and Geddes, J.W. (2006). Synaptic Mitochondria Are More Susceptible to Ca2+Overload than Nonsynaptic Mitochondria. Journal of Biological Chemistry 281, 11658–11668. 10.1074/jbc.M510303200.

102. Stiernagle, T. (2006). Maintenance of C. elegans: Transferring worms grown on NGM plates. In WormBook 10.1895/wormbook.1.101.1.

103. Evans, T. (2006). Transformation and microinjection. WormBook. 10.1895/wormbook.1.108.1.

104. Praitis, V., and Maduro, M.F. (2011). Transgenesis in C. elegans. In Methods in Cell Biology (Elsevier Inc.), pp. 159–185. 10.1016/B978-0-12-544172-8.00006-2.

105. Gong, J., Yuan, Y., Ward, A., Kang, L., Zhang, B., Wu, Z., Peng, J., Feng, Z., Liu, J., and Xu, X.Z.S. (2016). The C. elegans Taste Receptor Homolog LITE-1 Is a Photoreceptor. Cell 167, 1252–1263.e10. 10.1016/j.cell.2016.10.053.

106. Doser, R., Knight, K.M., Deihl, E., and Hoerndli, F. (2023). Subcellular Imaging of Neuronal Calcium Handling *In Vivo* Journal of Visualized Experiments. 10.3791/64928.

