## Supplemental material and figures for "Activity-dependent Mitochondrial ROS Signaling Regulates Recruitment of Glutamate Receptors to Synapses"

SUPPLEMENTAL METHODS

Imaging of Cytoplasmic GCaMP6f

The AVA neurite was located and continuous autofocus set as described above. Then, a 60 s image stream was collected with a 488 nm imaging laser (set to 0.1% power and an attenuation of 10) and a 250 ms exposure. Localized ChRimson activation (see the Methods section “Localized ChRimson Activation”) was triggered every 30 seconds using MetaMorph’s “Trigger Components” feature starting 15 seconds after the start of the stream acquisition. Imaging of GCaMP6f fluorescence was continuous throughout the optical activation protocol.

SUPPLEMENTAL TABLES

### **Table S1: List of Strains Used in This Study**

| Strain | Genotype | Contents |
| --- | --- | --- |
| FJH 18 | *akIs141* II; *glr-1(ky176)* III | *Prig-3::GLR-1::GFP* (integrated) |
| FJH 402 | *lin-15(n765ts), lite-1(ok530)* X; *csfEx160* | pRD30 + pRD15 + pJM23 + pCT61 |
| FJH 412 | *lin-15(n765ts), lite-1(ok530)* X; *csfEx167* | pRD30 + pAS1 + pJM23 |
| FJH 416 | *lin-15(n765ts), lite-1(ok530)* X; *csfEx168* | pRD36 + pRD15 + pJM23 |
| FJH 555 | *akIs141* II*; glr-1(ky176)* III; *csfEx188* | *akIs141;* pRD36 + pJM23 + pCT61 |
| FJH 582 | *lin-15(n765ts)* X; *csfEx210* | pRD36 + pDM1442 + pJM23 |
| FJH 635 | *lin-15(n765ts)* X; *csfEx234* | pDM1442 + pJM23 + pCT61 |
| FJH 641 | *lin-15(n765ts)* X; *mcu-1(ju1154)* IV; *csfEx261* | pRD30 + pAS1 + pJM23 +pCT61 |
| FJH 644 | *lin-15(n765ts), lite-1(ok530)* X; *csfEx264* | pKK01 + pRD30 + pJM23 + pCT61 |
| FJH 647 | *lin-15(n765ts), lite-1(ok530)* X*; mcu-1(ju1154)* IV; *csfEx264* | pKK01 + pRD30 + pJM23 + pCT61 |
| FJH 690 | *glr-1(ky176) III; csfEx268* | pKK?? + pDM1442 + pCT61 |

### **Table S2: List of Plasmids**

| **Name** | **Composition** | **Source** |
| --- | --- | --- |
| pRD15 | *Pflp-18::TOMM-20::roGFP:: let-858* 5’UTR | Hoerndli Lab, Colorado State University |
| pRD30 | *Pflp-18::ChRimson::tdTomato::let-858* 5’UTR | Hoerndli Lab, Colorado State University |
| pRD36 | *Pflp-18::TOMM-20::KillerRed::let-858* 5’UTR | Hoerndli Lab, Colorado State University |
| pKK01 | *Pflp-18::mito4x-GCaMP6f::let-858* 5’UTR | Hoerndli Lab, Colorado State University |
| pKK07 | *Pflp-18::TOMM-20::tdTomato::let-858* 5’UTR | Hoerndli Lab, Colorado State University |
| pED01 | *Pflp18::ChRimson::mCherry* | Hoerndli Lab, Colorado State University |
| pAS1 | *Prig-3::GCaMP6f::unc-54* 5’UTR | Stetak Lab, University of Zurich |
| pDM1442 | *Prig-3::SEP::GLR-1::unc-54* 5’UTR | Maricq Lab, University of Utah |
| pJM23 | *Plin-15::lin-15^+^* | Maricq Lab, University of Utah |
| pBSKS | Filler DNA | Stratagene |
| pCT61 | *Pegl-20::nls::DsRed* | Hoerndli Lab, Colorado State University |

SUPPLEMENTAL FIGURE 1

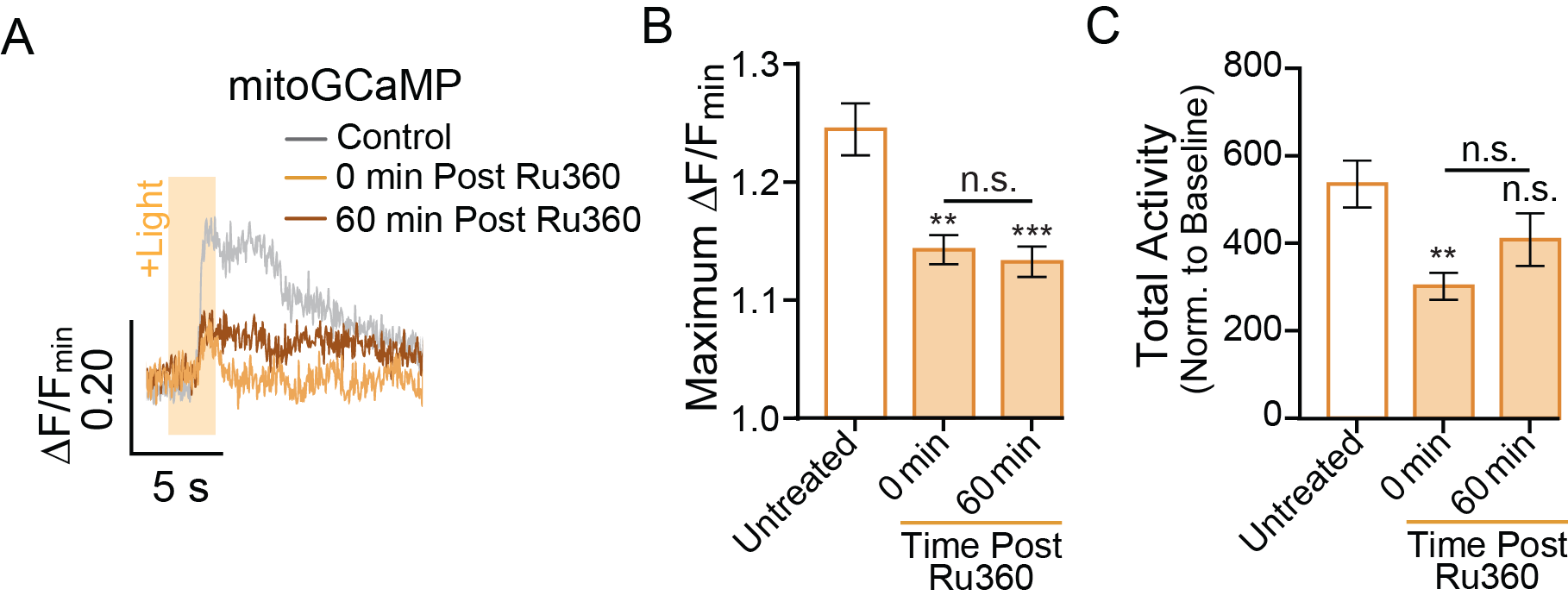

**Supplemental Figure 1:** A) The change in mitoGCaMP fluorescence (normalized to the minimum fluorescence for the region of interest; ∆F/F_min_) following optical stimulation (+Light) of the AVA neuron in untreated controls as well as Ru360 treated worms at 0- or 60-minutes post treatment. B) Quantification of the average maximum ∆F/F_min_ of mitoGCaMP during a 2.5-minute recording during which the AVA neuron was optically activated every 30 seconds (n≥20 mitochondria from 4-5 animals per group). C) Total mitoGCaMP activity (normalized to baseline fluorescence) over an entire 2.5-minute recording (calculated from same dataset as in panel B). Data is represented as mean ± s.e.m.; n.s. = not significant, **: p<0.005, ***: p<0.0005 compared to controls or the indicated experimental group using a one-way ANOVA with a Dunnett’s test.

SUPPLEMENTAL FIGURE 2

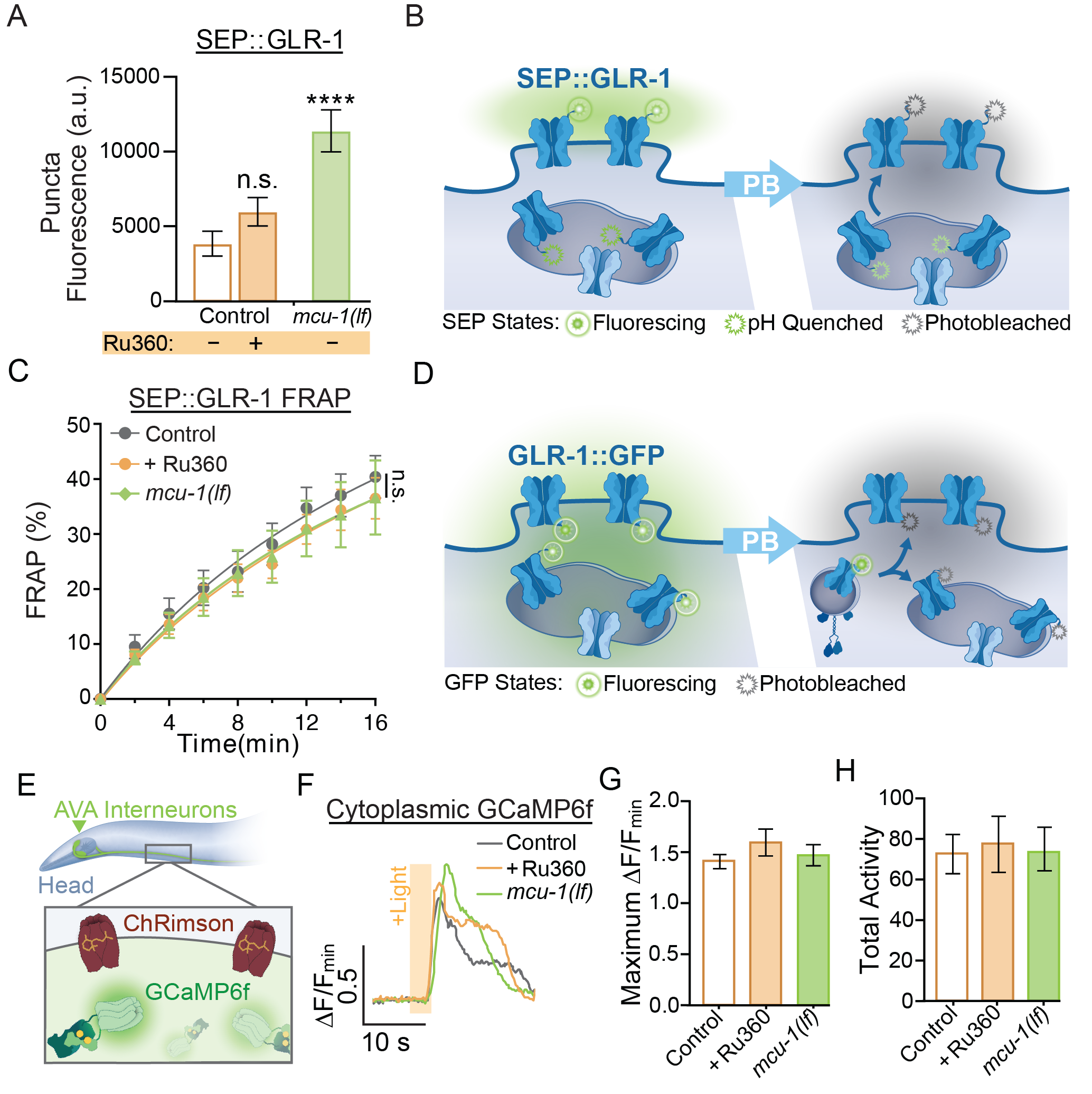

**Supplemental Figure 2:** A) SEP puncta fluorescence (a.u. = arbitrary units) in control, Ru360 treated, and *mcu-1(lf)* animals prior to FRAP (n≥8 animals per group). B) Illustration of SEP::GLR-1 localization. Following photobleaching (PB) of fluorescing SEP (attached to GLR-1 positioned at the synaptic membrane), recovery of SEP fluorescence is indicative of exocytosis of GLR-1 from transport vesicles or synaptic endosomes. C) Percent of SEP recovery after photobleaching (FRAP) over 16 minutes post photobleaching in each experimental group (n≥8 animals per group) from the same dataset as Figure 2C. n.s = not significant as determined by comparing the fitted curves using an extra sum-of-squares F-test with Bonferroni correction. D) Illustration depicting the localization of GLR-1::GFP to the synaptic membrane or in endosomes. Following PB of GFP, the fluorescence recovery indicates that new GLR-1 has been transported and delivered to the synaptic membrane or endosome within the region of interest (ROI). E) Illustration of reagents used for optical stimulation during imaging of GCaMP6f. ChRimson was localized to the plasma membrane and GCaMP6f was present in the cytosol in AVA interneurons. F) ∆F/F_min_ of GCaMP6f following optical stimulation. G) Maximum ∆F/F_min_ of GCaMP6f during a 1.5-minute recording during which the AVA was optically activated every 30 s (n≥10 animals per group). H) Total GCaMP6f activity (normalized to baseline fluorescence) from the same dataset as in (G) during the entire 1.5-minute recording. Data is represented as mean ± s.e.m.; n.s = not significant, ****: p<0.0001 compared to controls or indicated experimental group using a one-way ANOVA with a Dunnett’s test.

SUPPLEMENTAL FIGURE 3

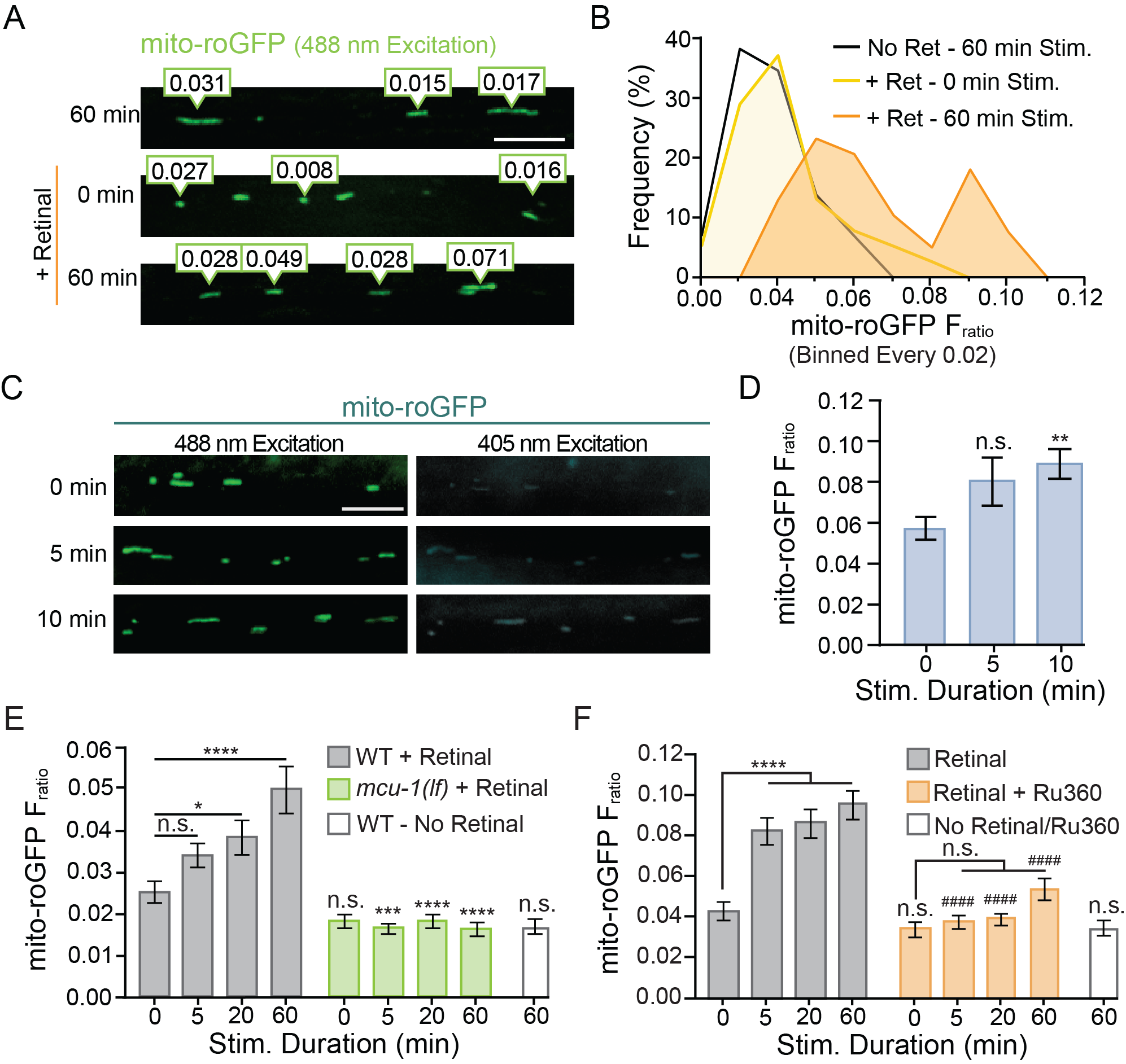

### **Supplemental Figure 3:** A) Representative image of the diversity observed in mito-roGFP F_ratios_ (emission due to 405 nm excitation / emission due to 488 nm excitation) in a portion of the AVA following repetitive optical activation. B) The frequency of observed mito-roGFP F_ratios_ expressed as a percent of the total observed F_ratios_ (binned every 0.02) from the same dataset as Figure 3C. C) Representative images of mito-roGFP fluorescence in a single z-plane when excited with 488 nm or 405 nm light following mechanosensory stimulation. D) Mito-roGFP F_ratio_ following 0, 5 or 10 minutes of repetitive mechano-stimulation (n>50 mitochondria from 8 animals per

### condition). E) The complete dataset from Figure 3E showing mito-roGFP F_ratios_ (Ex405/Ex488)

### observed following 0, 5, 20 or 60 minutes of repetitive optical stimulation in controls and *mcu-1(lf)* (n>34 mitochondria from 8 animals per group). F) The complete dataset from Figure 3G mito-roGFP F_ratios_ observed at 0, 5, 20 or 60 minutes of repetitive light stimulation in Ru360 treated animals compared to controls (n>38 mitochondria from 8 animals per group). Data is represented as mean ± s.e.m.; n.s = not significant, **: p<0.005, ***: P<0.0005, ****: p<0.0001 compared to non-activated controls or indicated experimental group using a one-way ANOVA with a Dunnett’s test.

### **SUPPLEMENTAL FIGURE 4**

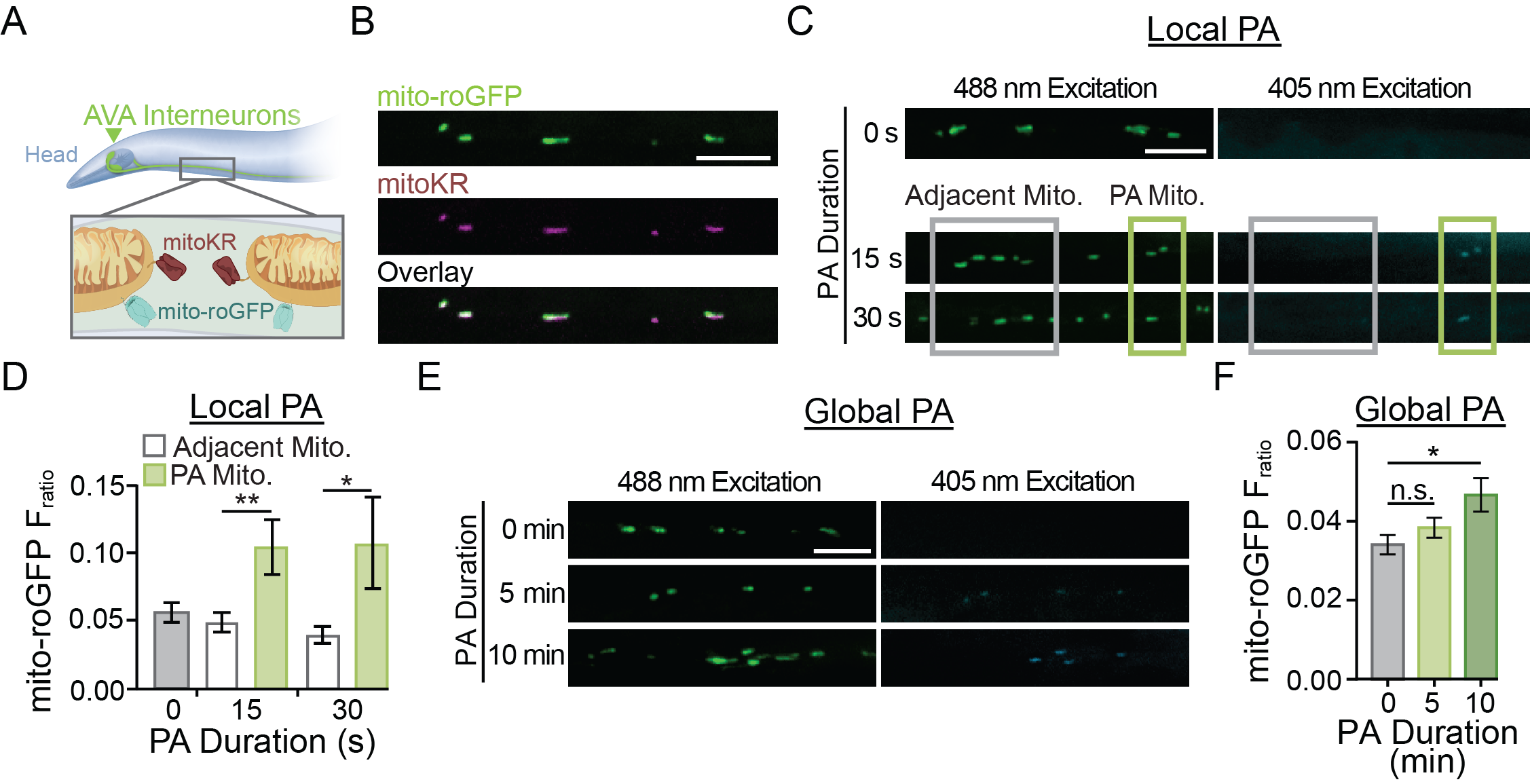

**Supplemental Figure 4:** A) Illustration depicting subcellular localization of mitoKR and mito-roGFP within the AVA neurites. B) Representative fluorescent images demonstrating the co-localization of mitoKR and mito-roGFP within the AVA neurite. C) Representative fluorescent images of mito-roGFP when excited by 405 or 488 nm light following 0, 15 or 30 seconds of PA directed at 1-3 mitochondria (green box). Localization of PA was considered to be spatially specific enough that neighboring mitochondria (grey box) were not exposed to the PA stimulus. D) The mito-roGFP F_ratio_ in mitochondria that were (green bars; n = 8 mitochondria from 8 worms per group) or were not (white bars; n > 20 mitochondria from 8 worms) targeted for PA as well as in worms without any additional optical activation (gray bars). n > 20 mitochondria from 8 worms per group; *: p<0.05, **: p < 0.005 using a paired t-test. No significant difference between the no light controls and the neighboring mitochondria (One-way ANOVA with Dunnett’s test). E) Representative fluorescent images of mito-roGFP excited by 488 or 405 nm light with 0, 5, or 10 minutes of consistent light (567 nm; 0.025mW/mm^2^) for global photoactivation (PA) of mitoKR. F) Quantification of mito-roGFP fluorescence ratio (F_ratio_, Ex405/Ex488nm) for each group (n>32 mitochondria from 8 animals per group). All scale bars = 5 µm. Data is represented as mean ± s.e.m.; *: p<0.05, n.s.= not significant using a one-way ANOVA with a Dunnett’s test.

### **SUPPLEMENTAL FIGURE 5**

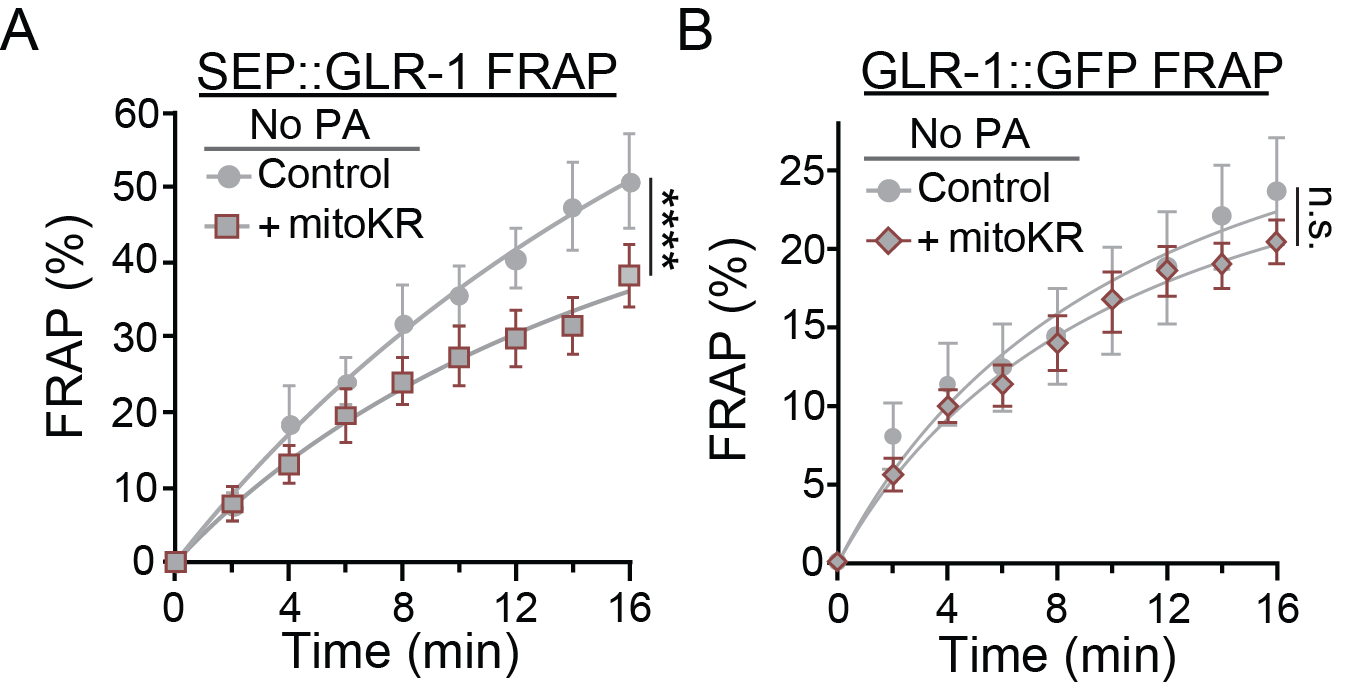

**Supplemental Figure 5:** A and B) Percent SEP (A) or GFP (B) fluorescence recovery after photobleaching (FRAP) over 16 minutes post-photobleaching and without photoactivation in controls or worms expressing mitoKR in the AVA neurons (n≥7 animals per group). Data is represented as mean ± s.e.m.; ****: p<0.0001, n.s. = not significant as determined by comparing the fitted curves using an extra sum-of-squares F-test with a Bonferroni’s correction for multiple comparisons.

**SUPPLEMENTAL FIGURE 6**

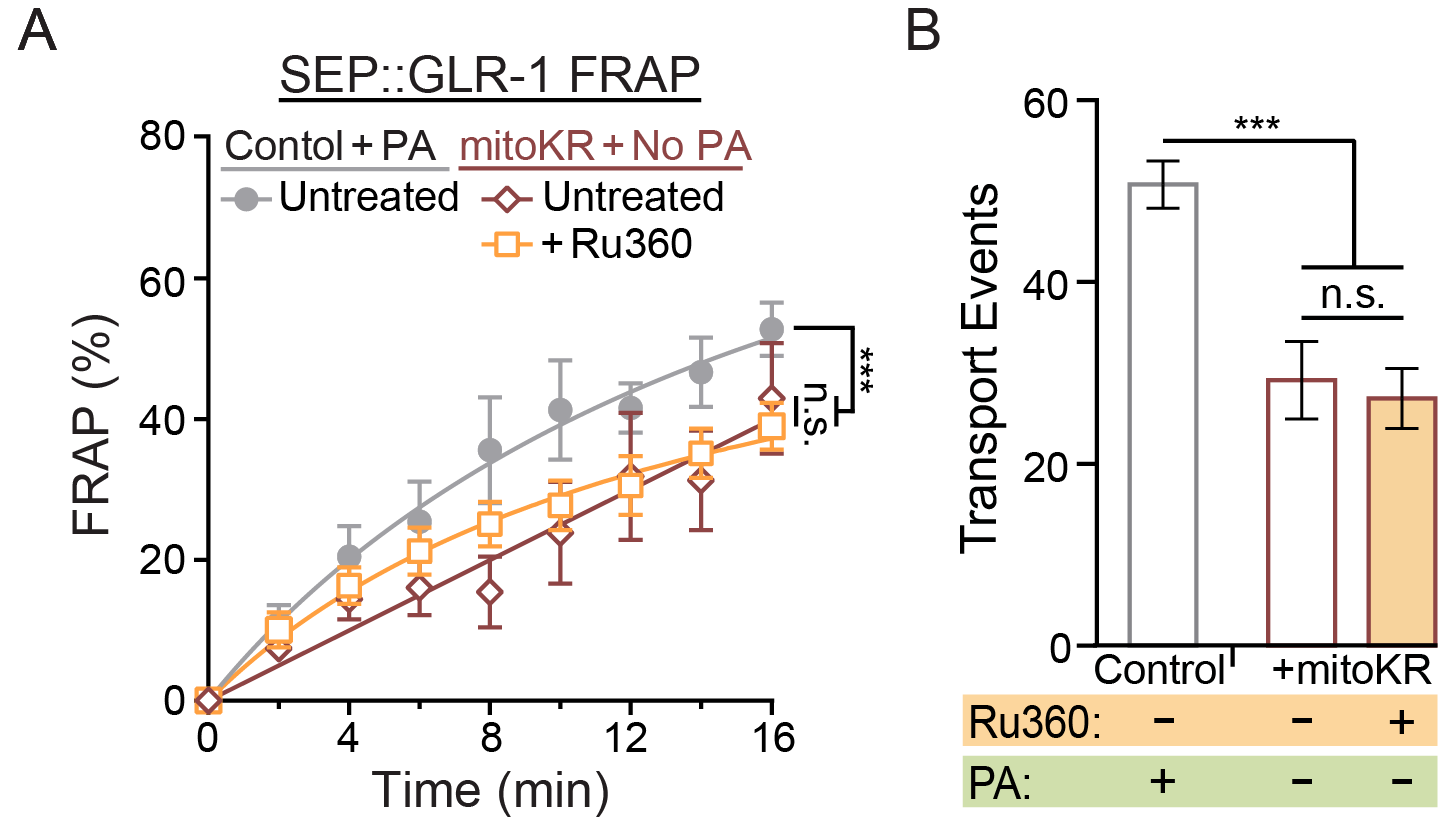

**Supplemental Figure 6:** A) Percent of SEP fluorescence recovery after photobleaching (FRAP) in the same experimental groups as (A) (n≥5 animals per group). B) Number of GLR-1 transport events quantified from 50-second-long kymographs in controls (n=10 animals per group) with local photoactivation (PA) as well as mitoKR-expressing worms with or without Ru360 treatment but without PA (n=5 animals per group). Data is represented as mean ± s.e.m.; n.s. = not significant, ***: p<0.0005 compared to controls using a one-way ANOVA with a Dunnett’s test.
